# Comparative genomics of the unusual arachnid order Solifugae spotlight the molecular and genetic basis for adaptations to arid habitats

**DOI:** 10.64898/2026.06.25.734573

**Authors:** Erika L. Garcia, Siddharth S. Kulkarni, Matthew R. Graham, Carlos Santibañez-Lopez, Prashant P. Sharma

**Affiliations:** Department of Integrative Biology, University of Wisconsin–Madison, Madison, WI 53706, USA; CSIR-Center for Cellular and Molecular Biology, Hyderabad, Telangana 500 048, INDIA; Academy of Scientific and Innovative Research (AcSIR), Ghaziabad, India; Department of Biology, Eastern Connecticut State University, Willimantic, CT 06226, USA; Department of Biology, Western Connecticut State University, Danbury, CT 06810, USA

**Keywords:** Comparative genomics, Desert adaptation, Heat shock proteins, Positive selection, Arachnida, solifuges

## Abstract

The evolutionary transition to terrestrial life required overcoming several physiological hurdles; however, such challenges were amplified in desert environments. While several xeric-adapted arachnids utilize permanent burrows or “sit-and-wait” foraging strategies as possible energy conservation adaptations in harsh habitats, camel spiders exhibit a counterintuitive, high-energy lifestyle. To investigate the molecular underpinnings distinguishing Solifugae within Chelicerata, we utilized a comparative genomics framework that incorporates a newly sequenced, previously unpublished solifuge genome. We identified lineage-specific expanded orthogroups and evaluated selective pressures acting upon paralogous sequences within our ingroup solifuge species. Additionally, we also focused on fatty acid-associated proteins and heat shock proteins to elucidate how Solifugae may have evolved such anomalous behaviors compared to their arachnid relatives. Our analyses revealed significant signatures of positive selection within key gene families across the solifuge lineage. Notably, paralogs within the cytochrome P450 and biotinidase families showed consistent evidence of selection across all three taxa, suggesting specialized metabolic or detoxification requirements. Furthermore, we identified candidate loci implicated in axonal guidance and lipid metabolism, and a specialized fatty acid enzyme repertoire. While subsequent research is required to determine whether some of the genomic signatures unveiled here are shared across a broader phylogenetic distribution within Solifugae, we establish a critical baseline for future functional validation.

## Introduction

Transitions to terrestrial life require organisms to overcome several major physiological challenges, including respiration, osmoregulation, thermoregulation, and exposure to environmental stressors such as desiccation and rapid temperature fluctuations (Little, 1990; Dunlop et al., 2013; Lozano-Fernandez et al., 2016; Perez & Aron, 2020; Benoit et al., 2023; Frumkin & Chipman, 2023). In terrestrial arthropods, the evolution of tracheal systems for respiration on land and various structural changes to the morphology of exoskeletons are some of the adaptations that enabled organisms to persist and thrive on land (Blomquist & Bagnères, 2010; Cloudsley-Thompson, 2012; Dunlop et al., 2013; Gefen et al., 2015; Tihelka et al., 2022; Wang et al., 2022; Frumkin & Chipman, 2023). In desert environments however, environmental stressors like solar radiation, extreme temperatures, and aridity imposed on organisms are intensified. Lineages that successfully underwent terrestrialization likely established the ancestral physiological framework that was subsequently refined to enable adaptation to arid conditions. Such taxa evolved unique morphological, physiological, and behavioral adaptations to survive extreme conditions of drought and heat.

One of the notable adaptive strategies of desert organisms is the ability to withstand long periods without nutrients, especially while under thermal stress. To mitigate the detrimental cellular damage inflicted by the exposure to harsh temperatures, xeric taxa have also evolved broad physiological mechanisms to reduce metabolic exertion under stressful states to avoid excess energy depletion (Klok et al., 2004; Perez & Aron, 2020). Other physiological responses to stress stimuli involve the synthesis and use of heat shock proteins (hsps). These proteins are the first line of defense, protecting against cellular degradation and ensuring proper cellular function, and thus are crucial for life to persist in adverse environmental conditions (Pockley, 2003; Haslbeck & Vierling, 2015; Z. Q. Miao et al., 2020; Voth & Jakob, 2017). Among ectotherms, thermoregulation is directly affected by their surrounding environment and must possess machinery to be able to counter detrimental thermodynamic influences on intracellular processes crucial for survival. Insights into the molecular evolution of desert taxa is necessary to understand patterns of adaptation in xeric environments–an assessment needed to predict the effects of ongoing and future climate change in some of the harshest habitats in the world.

In addition to thermal stress endured in deserts habitats, the physiological mechanisms responsible for osmoregulation in harsh climates have evolved to allow life to persist. For example, among some terrestrial arthropods, strategies to endure life in arid environments may include the suppression of metabolic rate during dehydration or stress, which in turn, reduces water loss by means of respiration (Addo-Bediako et al., 2001; Marron et al., 2003; White et al., 2007; Oladipupo et al., 2022). Cuticular evaporative loss, however, is suggested to be the leading cause of desiccation in insects (Zachariassen, 1996; Chown and Nicolson, 2004; Benoit, 2009; Chown et al. 2011), and the cuticular permeability of the exoskeleton depends on the structural composition of the hydrophobic layer (Blomquist & Bagnères, 2010; Gefen et al., 2015). For example, the length of carbon chains in the cuticular hydrocarbons, the alkanes profile, and the nature of the fatty acid chains (unsaturated versus saturated), are several molecular profiles that influence desiccation resistance (Blomquist & Bagnères, 2010; Gefen et al., 2015; Wang et al., 2022). Additionally, other osmoregulatory adaptations may include the activation of voltage-gated ion channels, such as in the Malphighian tubule excretory system which triggers a water recovery operation through osmosis for functional re-use (Ramsay, 1964; O’Donnell & Machin, 1988; Beaven et al., 2024). Although physiological adaptations to arid environments have been thoroughly studied in different insect groups (Addo-Bediako et al., 2001; Chown et al., 2002; Duncan & Dickman, 2009; Guillén et al., 2015; Rane et al., 2019; Holmes & Benoit, 2019; Perez & Aron, 2020; Benoit et al., 2023; Wang et al., 2023; Kumar et al., 2024; Roberts et al., 2025), insights into the molecular evolution associated with such physiological adaptations in arachnids remains limited.

The arachnid order Solifugae is among the smaller arachnid groups with a near global distribution, primarily inhabiting warm humid, semidesert, and desert environments (Cloudsley-Thompson, 1977; Brookhart & Brookhart, 2006; Hebets et al., 2024). Solifuges, compared to other arachnids adapted to such environments, are uniquely recognized for their fast running speeds (Punzo, 1998a), high metabolism (Lighton & Fielden, 1996), and ability to withstand extreme heat in some of the hottest places on the planet (Mildrexler et al., 2006; Brookhart & Brookhart, 2006; Hebets et al., 2024). Unlike spiders or some scorpions that build permanent burrows as an adaptive strategy for thermal regulation (Polis et al. 1986; Adams et al. 2016), solifuges are documented to have less efficient burrowing habits. Counterintuitive to energy conservation under stressful conditions, a new burrow is constructed after the end of an activity period (Gore & Cushing, 1980; Wharton, 1986) and appear to be constructed with extreme vigor (Muma, 1966a). Moreover, as many desert arachnids employ a “sit and wait” behavioral foraging strategy, solifuges are generally highly mobile, cursorial predators capable of covering large distances in a rapid, and seemingly random, manner (Muma, 1966b; Wharton, 1986; Punzo 1998a). Of the few studies that have documented periods of solifuge activity, such behavior may last for at least three hours (Hrušková-Martišová et al., 2010; Punzo 1998a). As mostly heat adapted animals, such counterintuitive behavior in comparison to other xeric adapted arachnids make solifuges an appealing test case among the chelicerates for studying and understanding desert adaptation. While recent studies revealed that there is a functional convergence on processes that drove arthropod terrestrialization (Lozano-Fernandez et al., 2016; Tihelka et al., 2022; Benítez-Álvarez et al., 2026), a focus on the molecular mechanisms underlying arid adaptations is non-existent in arachnids.

As a first step, we leverage several published chelicerate genomes and contribute one high-quality unpublished solifuge genome assembly to explore some of the molecular underpinnings enabling solifuges to thrive in xeric environments around the world (Mojave Desert, Central Asian deserts, and the Iberian Peninsula). In scrutinizing a subsample of chelicerate genomes, a group characterized by multiple independent colonization events throughout their evolutionary history (Sharma, 2017; Ballesteros & Sharma, 2019; Sharma & Gavish-Regev, 2025; Benítez-Álvarez et al., 2026), we identify unique and functionally enriched orthogroups specific to solifuge taxa. Our comparative genomic approach exposes the molecular machinery of desert survival, pinpointing the mechanisms driving adaptation to extreme aridity in this historically understudied arachnid group.

## Material and Methods

### Genome sequencing and annotation

An adult female specimen of *Hemerotrecha serrata* was collected at Afton Canyon Campground, CA (35.038°, −116.3823°) on March 13, 2023. The specimen was found active at approximately 7:50 AM (Graham). This specimen was shipped to the laboratory, and subsequently flash-frozen at −80 °C. Genomic DNA was extracted from the frozen chelicerae and a leg; two PacBio Sequel II libraries were prepared and sequenced across two SMRT cells at the Yale Center for Genomics.

### Assembly

PacBio HiFi CCS reads were assembled independently using hifiasm v.0.15.1-r329 (H. Cheng et al., 2021) assemblers to generate contigs. The assembly graph generated by hifiasm was converted to a set of primary contigs in multi-fasta format using “awk ’/^S/{print “>“$2;print $3}’”. Genome length and N50 were calculated using assemblyStatistics v.1.1.3 (Lin, 2023). In order to quantify biological completeness of our contig set, we used the package BUSCO v.5.4.7 (Manni et al., 2021) with the arthropod_odb10 ancestral lineage data set.

### Annotation

A library of repetitive elements was compiled using RepeatModeler2 v.2.0.5 (Flynn et al., 2020) for the genome assembly. The identified repeats were soft masked using RepeatMasker v.4.1.5 (Smit et al., 2015). For annotation, RNASeq data was not available for *Hemerotrecha*, annotation was performed by mapping the masked genome to Arthropod OrthoDB (Kuznetsov et al., 2023). Full gene structure annotations were predicted using BRAKER3 v.3.0.3 (Gabriel et al., 2024).

### Phylogenetic Reconstruction and Divergence Time Estimation

To explore gene family evolution, we used the genome projects of three solifuge (*H. serrata*, *Paragaleodes pallidus* and *Gluvia dorsalis*) alongside 11 additional chelicerate genomes (one sea spider, and 10 arachnids; Supporting Information 1). Orthologous sequences were identified from the proteomes of these 14 taxa with BUSCO v.5. A custom script was used to extract BUSCO sequences in the arthropod odb10 lineage gene set and sequences were aligned using mafft v.7. Sequences were trimmed using GBlocks v.0.91b with a 10-site minimum block length (*b4 = 10*) and permitting no sites with gaps (*b5=n*). Alignments with a minimum of 10 taxa were retained, resulting in 1,008 alignments. Tree topology was inferred with IQTree v.3 (Wong et al., 2026) with automated model fitting for each partition *(-m = MFP*) and 1000 ultrafast bootstrap replicates to infer nodal support (*-bb 1000*).

Divergence times were estimated using MCMCtree v4.9 and CODEML v4.9 (within the PAML suite; Yang 2007) based on the 1,008-locus alignment and the constrained ML phylogeny. We applied four temporal constraints derived from established benchmarks to the topology (Supporting Information 1). To optimize computational efficiency, we employed the approximate likelihood method, calculating a Hessian matrix in CODEML (dos Reis & Yang, 2011; 2019) using empirical base frequencies and a WAG+G6 substitution model. Posterior distributions were estimated under a correlated molecular clock model across four independent MCMC chains. Each chain was executed for 12 million generations, with the initial 40,000-generation discarded as burn-in to ensure convergence.

### Gene Family Evolution

Orthologous groups were identified by retrieving the longest isoforms from the 14 proteomes using a custom Python script and *seqtk* (https://github.com/lh3/seqtk), followed by analysis in OrthoFinder v2.5.4 (Emms & Kelly, 2019). Functional annotation was performed using eggNOG-mapper v2.1.12 (Huerta-Cepas et al., 2019), using the longest sequence per orthogroup as a representative. To minimize phylogenetic noise from lineage-specific repetitive elements, genes annotated as transposable elements (TEs) were removed, and orthology inference was re-executed on the filtered dataset. Furthermore, we excluded Hierarchical Orthologous Groups (HOGs) with high gene copy number variance (>580) and those present in only a single species. Gene family expansions and contractions were modeled using CAFE5 v1.1 (Mendes et al., 2021). We evaluated five evolutionary models: Uniform and Poisson distributions for the root state, as well as configurations incorporating two gamma rate categories. To account for potential assembly or annotation artifacts, an error model was estimated under both root Uniform and root Poisson configurations. Significant expansions and contractions within Solifugae and individual species were identified using a significance threshold of p < 0.01.

### Test for positive selection in expanded orthogroups

To test for positive selection across expanded gene families per ingroup within solifuges, we isolated amino acid sequences pertaining to each respective expanded gene family expanded significantly (p < 0.01) using *seqtk* the program subseq from the suite *seqtk* then and aligned each FASTA file them using MAFFT (using --auto) (Katoh & Standley, 2013). Next, we trimmed our aligned sequences using TrimAI (Capella-Gutiérrez et al., 2009) using the 0.75 gap-threshold. To generate codon alignments for downstream positive selection analyses, we translated our amino acid sequences to nucleotide sequences using *backtranseq* in the EMOSS package (Rice et al., 2000). We subsequently used both the amino acid sequence files and associated nucleotide files to generate final codon alignments using *pal2nal.pl* (Suyama et al., 2006). For each expanded family in each solifuge species, we reconstructed individual Maximum Likelihood trees in IQTree v.3 (Wong et al., 2026) using the -m MFP+MERGE and -B 1000 option.

Using the expanded orthogroups unique to the solifuge lineage, we investigated whether specific gene copies within those expanded orthogroups showed signs of an adaptive advantage. First, we isolated each expanded orthogroup by individual solifuge species and selected those orthogroups with more than five paralogs for analysis. Next, we conducted a preliminary scan of our codon alignments and corresponding tree files for possible sources of error, such as uninformative branch lengths or duplicate sequences, that would impact downstream positive selection analyses. To do this, we used the *HyPhy v 2.9.96* software package and employed the BUSTED-E model to eliminate residual alignment errors by applying an “error-sink” stringent filtering method (Selberg et al., 2025). Resulting filtered codon alignments were then used as input to test for episodic positive selection across all branches for each orthogroup using adaptive Branch-Site Random Effects Likelihood (abSREL; Kosakovsky Pond et al., 2011). This analysis incorporated synonymous rate variation (--srv Yes), and support for multiple nucleotide substitutions (Double + Triple), with no target branch pre-specified.

To identify specific loci were under episodic selection, we used a modified R script (; parse_significant_results.R) to parse aBSREL results, identifying the branches and nodes with an estimated Holm-Bonferroni uncorrected p-value of < 0.05. Of those with loci that were in line with episodic selection, we performed functional characterization using InterProScan, to predict protein domains and assign putative Gene Ontology (GO) terms (Jones et al., 2014). GO term annotations were then iteratively retrieved using a custom R script, by selecting lowest E-scores for query matches for each unique transcript and using the QuickGO database (Binns et al., 2009). This custom script and the modified aBSREL parsing script are available at (Github link).

### Test for selection in orthogroups containing stress-immune and fatty acid metabolism proteins

Previous studies have shown that Heat Shock Proteins (HSPs) and genes involved in general metabolic pathways, including those related to fatty acid metabolism, may represent genomic adaptations to environmental stressors such as heat and desiccation (e.g., Carmel et al., 2011; Van Dooremalen et al., 2011; Haslbeck & Vierling, 2015; Advani et al., 2016; Chen et al., 2018; Wang et al., 2022; Wang et al., 2023). Therefore, we retrieved all HOGs functionally annotated as HSPs or as being involved in fatty acid metabolism, as identified through our EggNOG annotation pipeline. None of these HOGs were inferred by CAFE to be expanding or contracting, suggesting that these gene families are stable across arthropods. We then tested for signatures of selection within these HOGs using the same procedure as described above.

### Per-site selection pressure of positively selected paralogs and prediction of protein structures

We further analyzed the expanded orthogroups and fatty acid associated sequences showing evidence of positive selection using Fast Unconstrained Bayesian AppRoximation (FUBAR) to identify which codons were influenced by positive evolutionary pressure (Murrell et al., 2013). We performed this analysis with the same input FASTA files used to run aBSREL after conducting the “error-sink” filtering method. From the resulting FUBAR JSON files, we parsed each file to obtain those specific residues with posterior probabilities of > 0.90, an indicator of positive selection, using a custom script.

Mapping selected amino acid residues can reveal the specific functional roles that grant a protein a selective advantage in several different ways. First, specific amino acid residues cooperatively interact with one another to achieve a stable folded conformation (Ragupathi et al., 2026). Hydrophobic side chains are predominately located in the interior of the protein, while polar side chains are exposed to the solvent (Malleshappa Gowder et al., 2014), whose charged residues facilitates solubility and prevents heat denaturation (Barik, 2020). In the same regard, the existing hydrophobic amino acid residues at the core allows for tighter packing of the globular protein, therefore making it resistant to unfolding (Okada et al., 2010; Gromiha et al., 2013). These hydrophobic residues present at the interior of the tertiary structure are also a general characteristic that increase the catalytic power of the active sites of an enzyme (Bartlett et al., 2002). Moreover, residues in the C- or N-terminus of a protein may further dictate the stability of the protein by increasing the half-life by preventing a protein from rapidly degrading (Varshavsky, 1996; Varland et al., 2015; Weber et al., 2020), can promote effective cellular signaling often regulated by the

N-terminus (Varland et al., 2015), or could impact protein abundance by elevating expression based on the presence of certain residues present in the C-terminus (Weber et al., 2020). Therefore, we opted to generate 3D tertiary structures to identify potential hotspots for selection, such as a possible concentration in active sites, N-terminal, or C-terminal domains, and to determine if these residues are primarily located on the surface or core of the protein structure. AlphaFold3 via the AlphaFold Server (https://alphafoldserver.com/) was then used to model the 3D tertiary structure of each of our protein sequences under positive selective pressure for Solifugae. We selected the representative predictive model with the highest-ranking score based on quality and predicted accuracy (Abramson et al., 2024). Final 3D protein structures with annotated residues reflecting positive selection signal were rendered using PyMOL (The PyMOL Molecular Graphics System, Version 3.0 Schrödinger, LLC).

## Results

### Genome assembly and gene family dynamics

We generated a high-quality de novo genome assembly for the solifuge species, *Hemerotrecha serrata*, using PacBio HiFi long-read sequencing, the first such genomic resource for the family Eremobatidae. PacBio sequencing on two SMRTbells generated 3.2M sequences (length=17 Gbp; mean length=5313.9 ; N50=5773 bp) and 3M sequences (length=22.3 Gbp; mean length=7336.2 bp; N50=8137 bp) high-quality long reads, respectively. The hifiasm assembly consisted of 613 scaffolds with 688 Mbp length with sequences ranging 0.004 Mbp to 52.59 Mbp, N50 of 18.6 Mbp (n=12) and N90 of 3.4 Mbp (n=39).

BUSCO completeness scores with Arthropoda Orthology Database version 10 were 97.9% complete with 93.4% single copy and 4.5% duplicated orthologs (Figure 1). This makes the *H. serrata* datasets the most complete solifuge genome to date, despite being a scaffold-level assembly. The availability of three multiple annotated genomes with high BUSCO completeness (>90%) facilitates the first examinations of gene family evolution within this group of arthropods. The *H. serrata* genome assembly consisted of a total of 59.94% repetitive DNA, with interspersed repeats comprising 58.01%. The majority of those repeats were categorized as unclassified repeats, while Retroelements category was the second more represented in our RepeatMasker results, followed by DNA transposons (Supplemental Figure S1). The genome assembly was deposited on NCBI under BioProject XXXX.

**Figure 1:**
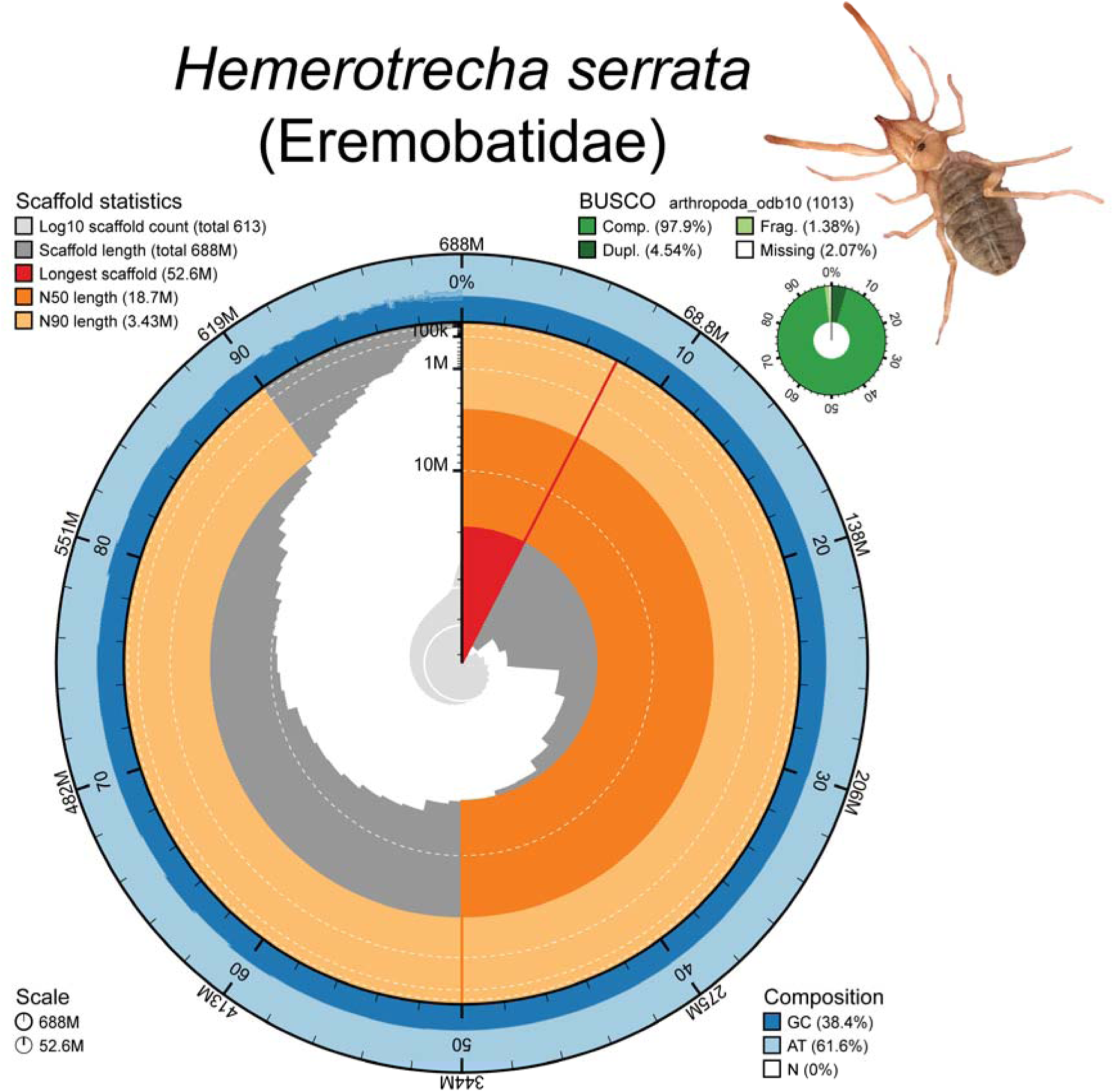
Snail plot summary of the assembly statistics and BUSCO scores for the *Hemerotrecha serrata* genome. The inner circular plot displays scaffold lengths in dark grey (arranged in descending order moving clockwise). The red radius represents the longest scaffold in the genome assembly. The pale grey area represents the log-transformed scaffold count. Dark and pale orange highlight the N50 and N90 sequence lengths, respectively. The outer ring displays the percentage distribution of GC (blue), AT (pale blue), and N (uncolored/missing) content mapped across corresponding bins. Assembly completeness is indicated by BUSCO scores (top right) assessed against the arthropod lineage-specific library. Image of the sequenced individual of *Hemerotrecha serrata*.

To investigate gene family dynamics in the branch subtending Solifugae, we next characterized gene family evolution across 14 chelicerate taxa, using OrthoFinder, which identified a total of 40,535 Hierarchical Orthologous Groups (HOGs; Figure 2A). Following stringent filtering to remove high-variance and lineage-specific HOGs, a dataset of 19,755 HOGs was retained for CAFE analysis, with results visualized on a fossil-calibrated phylogeny (Figure 2B). For temporal context, we built a time-calibrated phylogenetic tree to understand gene family evolution. The diversification of crown group of

**Figure 2.**
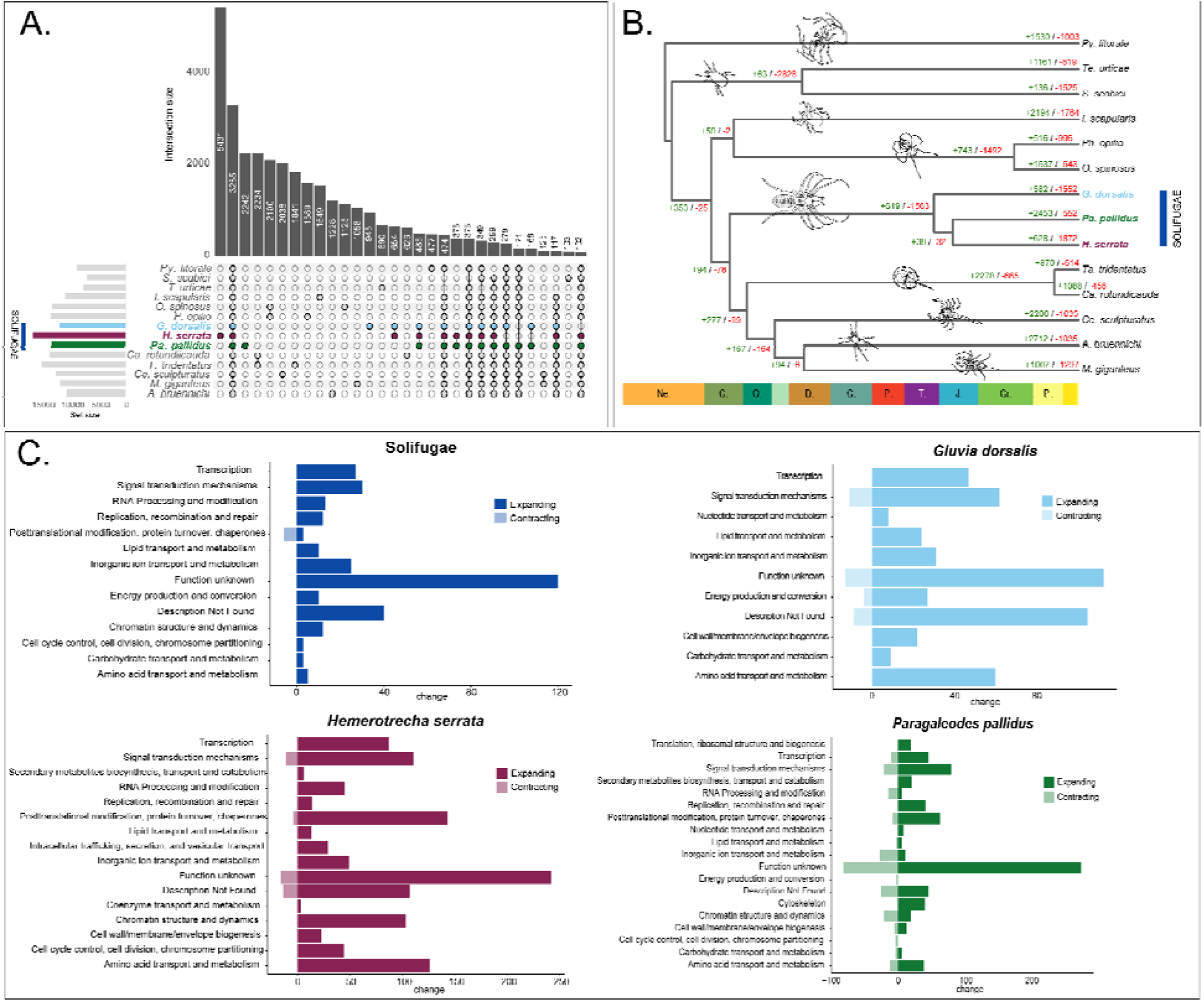
Comparative genomic and gene family evolution analysis across Solifugae and ingroup chelicerates. A.) Upset plot represents the number of HOGs across sampled chelicerates. Dark circles with connected circles indicate the number of shared HOGs between those intersected taxa. The horizontal bar chart on the left (Set size) represents the total number of orthogroups identified per species. Solifugae lineages (*Gluvia dorsalis* in light blue, *Hemerotrecha serrata* in burgundy, and *Paragaleodes pallidus* in green) are highlighted. B.) MCMCTree time-calibrated tree and gene family expansion/contraction rates across ingroup chelicerates. showing the evolutionary relationships among the sampled chelicerate species, with monophyletic Solifugae highlighted. Numbers at the nodes and terminal branches indicate the estimated number of expanded gene families in green (+) and contracted gene families in red (-). The chronostratigraphic scale at the bottom denotes geological periods. (C) Functional annotation of expanding and contracting gene families based on Clusters of Orthologous Groups (COG) categories for gene families undergoing significant size changes. Plots show the net change for the entire order Solifugae (top left, dark blue), as well as species-specific profiles for *Gluvia dorsalis* (light blue), *Hemerotrecha serrata* (burgundy), and *Paragaleodes pallidus* (green). Darker shades denote expanding families, while lighter, desaturated shades extending to the left of the zero-axis represent contracting families.

Solifugae was estimated within the Triassic, in agreement with previous studies (Kulkarni et al., 2023; Garcia et al., 2026). On this dated tree topology, we implemented a CAFE model that incorporated a Poisson distribution to estimate ancestral gene family sizes at the root, and a discrete gamma distribution (k = 2) to account for rate variation among different gene families (Pk2) since it minimized the negative log-likelihood value, and is the most parameter-rich model. Our results show that the MRCA of Solifugae experienced more contractions (-1563 HOGs) than expansions (+619; Figure 2C). Similar patterns were observed for *Gluvia dorsalis* (+582 / −1552) and *Hemerotrecha serrata* (+628 / −1872); but the opposite trend is observed in *Paragaleodes palidus* (+2453 / −552; Figure 2C). After applying statistical thresholds (Viterbi p < 0.05), we identified 229 unique HOGs undergoing significant expansion and 51 undergoing contraction across the solifuge lineage. Rather than being restricted to the deep ancestral node (MRCA), these figures capture cumulative evolutionary shifts across the entire ordinal topology, including species-specific terminal branches. Of these 229 expanded orthogroups, 61 (encompassing 331 individual gene copies) are expanding specifically at the root of Solifugae, while a single HOG comprising six gene copies undergoes contraction at this same deep node. Functional annotation revealed that most of these HOGs either fell into a Cluster Orthologous Gene (COG) category S (Function Unknown) or lacked a descriptive annotation entirely. In contrast, the contracted HOGs were primarily associated with posttranslational modification, protein turnover, and chaperones (COG category O). At the species level, significant expansions consistently outpaced contractions (Figure 2C). Consistent with the lineage-wide trends, uncharacterized proteins and sequences lacking functional descriptions constituted the majority of expanded HOGs within all three species.

### Evidence of Episodic Diversification and Per-Site Selection Signal

Out of the 43 total expanded HOGs recovered for Solifugae, we removed 11 unique orthogroups with fewer than five transcripts, leaving 29 HOGs for formal testing within each solifuge species. From those tested, 17 independent HOGs exhibited evidence for some degree of selection, many of which were supporting transcripts with strong positive episodic selection (Table 1). Our analyses supported directional selection for *H. serrata*, *G. dorsalis*, and *P. pallidus* yielding 16, 15, and 15 sequences, respectively.

**Table 1:**
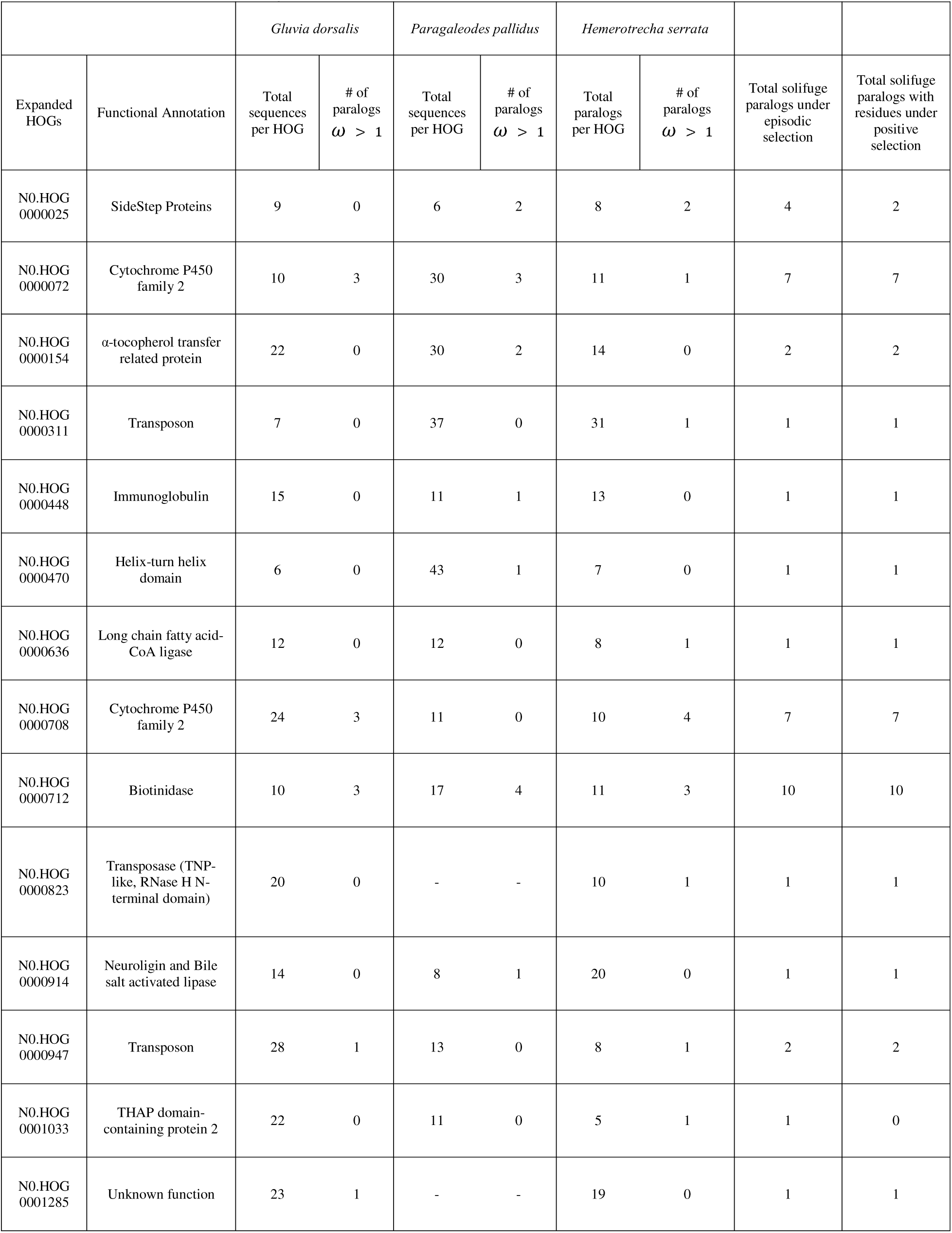
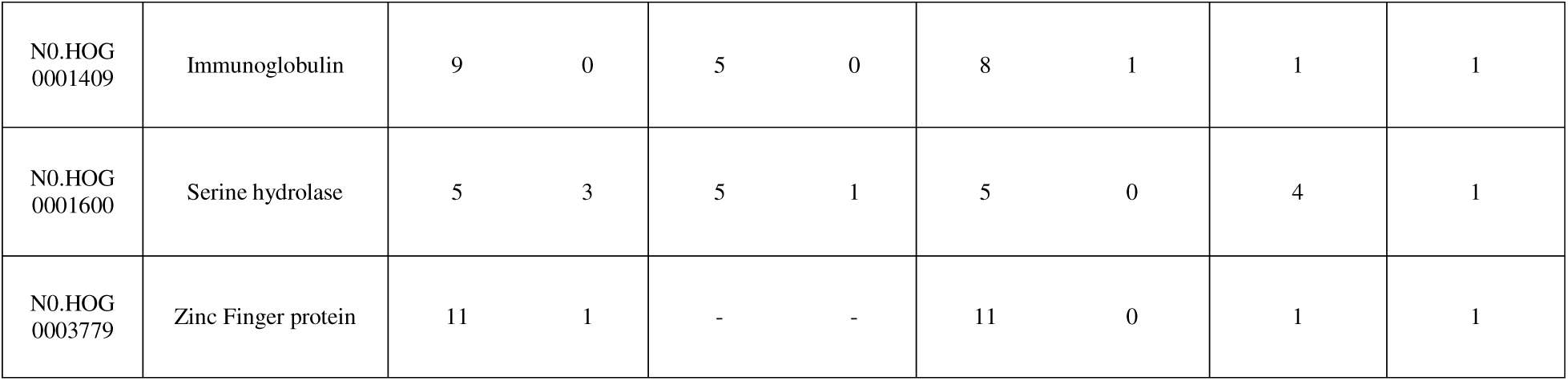
Summary of expanded Hierarchical Orthogroups (HOGs) within Solifugae showing the number of inparalogs with evidence of episodic selection (aBSREL, uncorrected p < 0.05) and number of inparalogs with amino acid residues under positive selection (FUBAR, PP > 0.90), along with their InterProScan functional annotations

InterProScan and subsequent GO term retrieval characterized the sequences under directional selection across a diverse functional landscape. Notably, all ingroup solifuges shared a core set of functional profiles, specifically matching Cytochrome P450 and Biotinidase/VNN families. Other categories, such as Sidestep proteins, α-Tocopherol transfer proteins, transposases, immunoglobulins, and serine hydrolases, were shared among some, but not all, lineages. Finally, several functional categories were species-specific, including a Zinc finger protein, neuroligin and bile salt-activated ligase, and a protein of unknown function. Full metadata for all positively selected paralogs, including omega (ω) values from aBSREL can be found in (Supplementary Information 1).

Of those 17 HOGs that were supported to show significant signal for positive episodic selection (Supporting Information 2), only 40 out of the 46 sequences tested across all three solifuge species supported at least one amino acid residue demonstrating positive selection signal (Supporting Information 3). Three sequences belonging to *Gluvia dorsalis* had the highest number of amino acid residues showing evidence for positive selection (PP >0.90) with a total of 72 residues pertaining to functionally annotated Biotinidase HOG (Table 1; Supplementary Information 2). The functionally annotated sequences for immunoglobulin (N0.HOG0001409: *Hemerotrecha serrata),* transposon (N0.HOG0000947:*Gluvia dorsalis*), and long-chain fatty acid-CoA ligase (N0.HOG0000636:*Paragaleodes pallidus*) only supported a single residue under adaptive selection. Of the 40 sequences tested, 30 those sequences showed dominant tendencies for hydrophobic residues, whereas a mere 7 sequences supported mostly hydrophilic residues under positive selection according to their hydrophobicity index at neutral pH (Monera et al. 1995). The remaining sequences had either equal residues showing affinities for hydrophobicity or hydrophilicity or were neutral (Supporting Information 3). The average length of each of the sequences examined was ∼931 with the minimum being 103 amino acids long and the maximum being 1000.

### Characterizing Heat-Shock and Fatty Acid Metabolism Pathways in Solifugae

While CAFE analysis primarily highlights dynamic shifts in gene family size, several canonical pathways associated with xeric adaptation – specifically those involving Heat Shock Proteins (HSPs) and fatty acid biosynthesis - exhibited patterns of copy-number stability rather than recent expansion (Viterbi p < 0.05). We identified 27 HOGs functionally annotated as Heat Shock Protein (HSP), nine of which were shared by, though not restricted to, the three solifuge species (Figure 5). Within this conserved core, one HOG (characterized as a small heat shock protein member of the HSP20 family) contained 12 paralogs in *P. pallidus*, eight in *H. serrata*, and six in *G. dorsalis*. This family also exhibited high baseline counts across the arachnopulmonates (>20 genes), and low counts in the remaining chelicerate lineages (>8). Similarly, a HOG annotated as a 70kda HSP cognate protein maintained high paralog counts across chelicerates. The remaining seven HOGs shared by solifuges showed consistent copy-number stability, although *P. pallidus* retained a slightly higher gene count across four of these groups.

In contrast, we identified 154 HOGs functionally annotated as involved in fatty acid metabolism, 29 of which were conserved across all three solifuge species. Notably, nine of these metabolic orthogroups were entirely restricted to the solifuge lineage. Within this order-specific subset, five HOGs maintained a single-copy orthology across the three taxa, suggesting a highly conserved, lineage-specific metabolic baseline. In the remaining four HOGs, *P. pallidus* and *G. dorsalis* exhibited a slightly greater number of inparalogs, whereas *H. serrata* maintained a single-copy state across all eight of these unique families. We further enriched the functional annotation of these HOGs using InterPro, uncovering five subfunctional classifications within the fatty acid metabolism gene families: (a) activation, (b) elongase, (c) metabolism, (d) reductase, and (e) synthase. Of the nine HOGs unique to solifugid species, six were assigned a putative elongase function and two a synthase function. Solifugae harboured more inparalog copies within HOGs with synthase function compared with the remaining chelicerates.

We tested for signatures of selection across 38 HOGs (29 fatty acid metabolism genes and nine HSPs) present in all three solifuge species (Figure 5). None of the paralog sequences assigned a metabolism function in solifuges showed evidence of significant positive selection (ω > 1, p > 0.05). In contrast, the majority of paralogs in the remaining functional categories showed significant positive selection (p < 0.05; Figure 5). Among the putative gene families unique to Solifugae, three exhibited significant signatures of positive selection across all three species, while three did not — all of the latter belonging to the elongase functional class. Additionally, one synthase-function gene family unique to Solifugae showed no significant evidence of positive selection (Figure 5). Regarding the HSP complement, one gene family showed significant positive selection across all three solifuge species, with two additional families showing species-restricted signatures — one in *G. dorsalis* and another in *H. serrata*.

Furthermore, we identified 154 five HOGs functionally characterized as enzymes protein involved in the metabolism of responsible for the elongation of very-long-chain fatty chains (VLCFA) acids that were restricted to the three solifuge species, from which 30 were present in all three solifugid species. Four of these orthogroups were strictly single-copy across Solifugae, indicating a highly conserved baseline for VLCFA synthesis within the order. However, the fifth HOG exhibited a lineage-specific expansion in *Gluvia dorsalis* (possessing four paralogs), while *Hemerotrecha serrata* and *Paragaleodes pallidus* maintained the ancestral single-copy state.

Families associated with Heat Shock Proteins (HSPs) and fatty acid biosynthesis were notably excluded from significant expansion results (Viterbi p < 0.05). From these, one HOG was functionally characterized as Heat Shock Protein A (HSPA), with *Pa. palidus* represented by two genes, *H. serrata* and *G. dorsalis* by one gene only. When compared to the other taxa studied, only the scorpion (*Centruroides sculpturatus*) and the horseshoe crab (*Carcinoscorpius rotundicauda*) had three genes.

## Discussion

### Gene family dynamics in an arid-adapted arachnid order

We characterized gene family evolution across Chelicerata using OrthoFinder and CAFE. In Solifugae, the overall genomic inventory was heavily dominated by net gene loss. Specifically, the most recent common ancestor (MRCA) of Solifugae experienced a total of 1,563 contracting families compared to only 619 expanding families. Interestingly, when applying strict statistical thresholds (Viterbi p < 0.05) to this ancestral node, the pattern of significance inverted: we identified 229 HOGs undergoing significant, targeted expansion, compared to just 51 driving the significant contractions. This broad trend of net background gene loss was maintained in the terminal taxa *Gluvia dorsalis* and *Hemerotrecha serrata* (Figure 2B). However, a stark exception was observed in *Paragaleodes pallidus*, which exhibited a markedly expansion-biased profile (Figure 2B) that warrants further research. Functionally, the contracted HOGs were significantly enriched for post-translational modification, protein turnover, and chaperone functions (COG category O)—a pattern consistent across all three solifuge species examined. The phylogenetic distances spanned by these three exemplars (which span the basal split of Solifugae; Kulkarni et al. 2023), together with the high BUSCO completeness of their assemblies, disfavor the interpretation that these contractions represent artifacts stemming from the incompleteness of their genomes. The predominance of contractions at deep nodes and most terminal branches, coupled with the overall low variation in gene content and genome size reported above, suggests a conserved and largely stable mode of genome evolution across Solifugae (Freitas & Nery, 2020; Rogers et al., 2022), with large-scale duplication events playing a limited role except in derived, lineage-specific bursts like that of *Paragaleodes*.

In total, we tested 43 expanded HOGs to determine if Solifugae shared parallel patterns of selection, and to elucidate which HOGs were providing lineage-specific advantages. From our analyses, we recovered 17 HOGs demonstrating strong support for episodic positive selection. Overall, only two HOGs showed at least one in-paralog under positive selection among all three ingroup solifuge species. Namely, N0.HOG0000072 that functionally annotated to the cytochrome P450 family 2 enzyme gene family (Table 1; Figure 3), and the other to biotinidase (N0.HOG0000072: Table 1; Figure 4). The remaining HOGs that demonstrated support for positive selection were either unique to the individual species or were consistent with one other species (Table 1). Nevertheless, our functional annotation suggests that these in-paralogs predominantly modulate cellular responses to oxidative stress, regulate locomotor behavior, control gene expression, or are broadly involved in metabolism.

**Figure 3.**
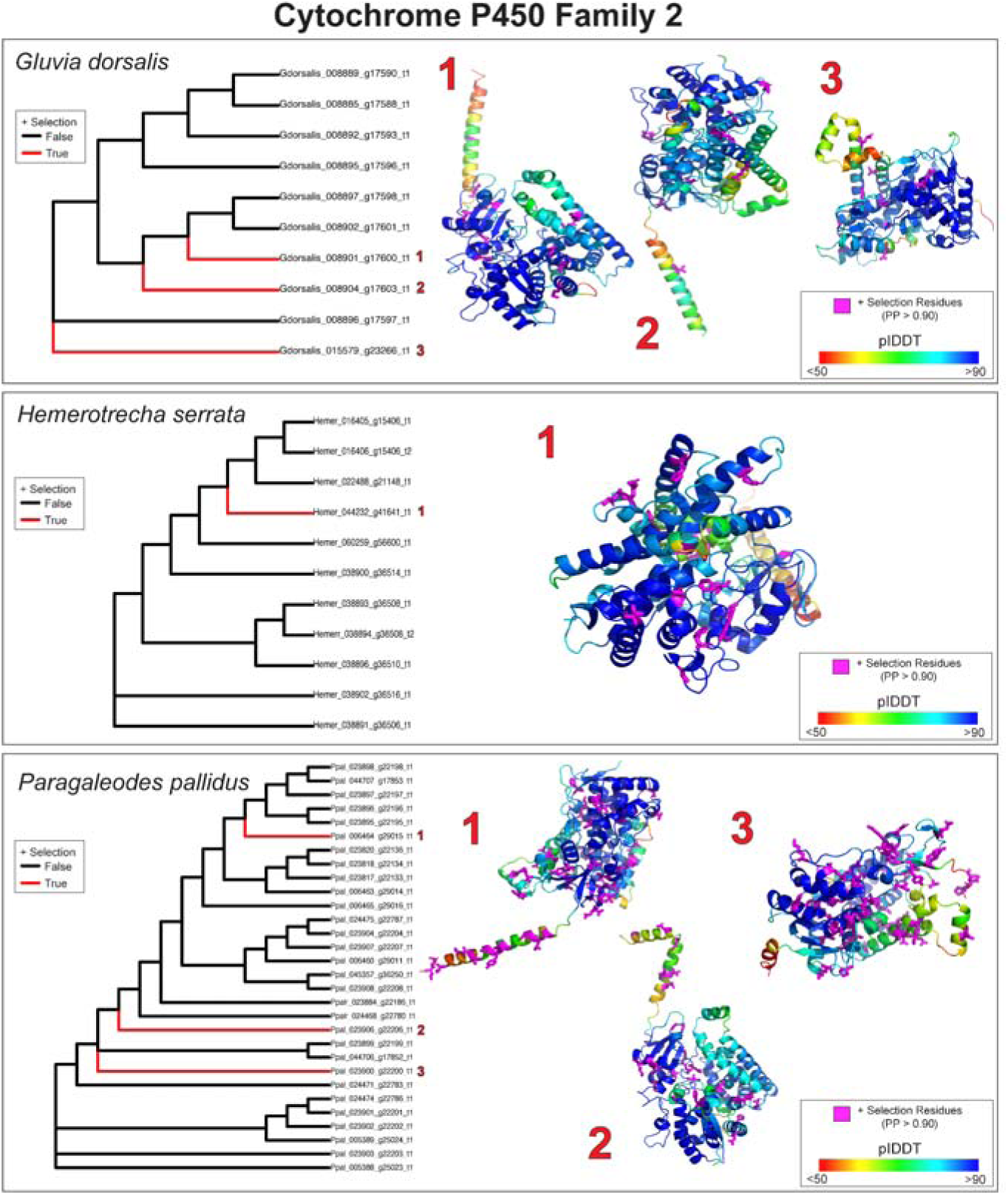
Inparalog positive selection results from aBSREL and corresponding 3D structural modeling of each respective positively selected protein for the functionally annotated cytochrome P450 family 2 protein (N0.HOG0000072) in solifuges. Maximum likelihood (ML) phylogenetic analysis and predicted 3D protein models for *Gluvia dorsalis* (top panel) and *Hemerotrecha serrata* (bottom panel). Left subplots display gene trees with all branches tested for episodic positive selection. Branches highlighted in red indicate inparalogs under significant positive selection (+ Selection (True) ; ω> 1, corrected p-value < 0.05), while black branches indicate no detected selection (False). Specific selected paralogs are numbered corresponding to their predicted structural models on the right. Structural models are colored by AlphaFold 3 per-residue confidence scores (pLDDT), ranging from very low confidence (red <50) to high confidence (blue >90). Pink coloration highlights the specific residues predicted to be under positive selection with a posterior probability (PP) > 0.90 recovered in FUBAR.

**Figure 4.**
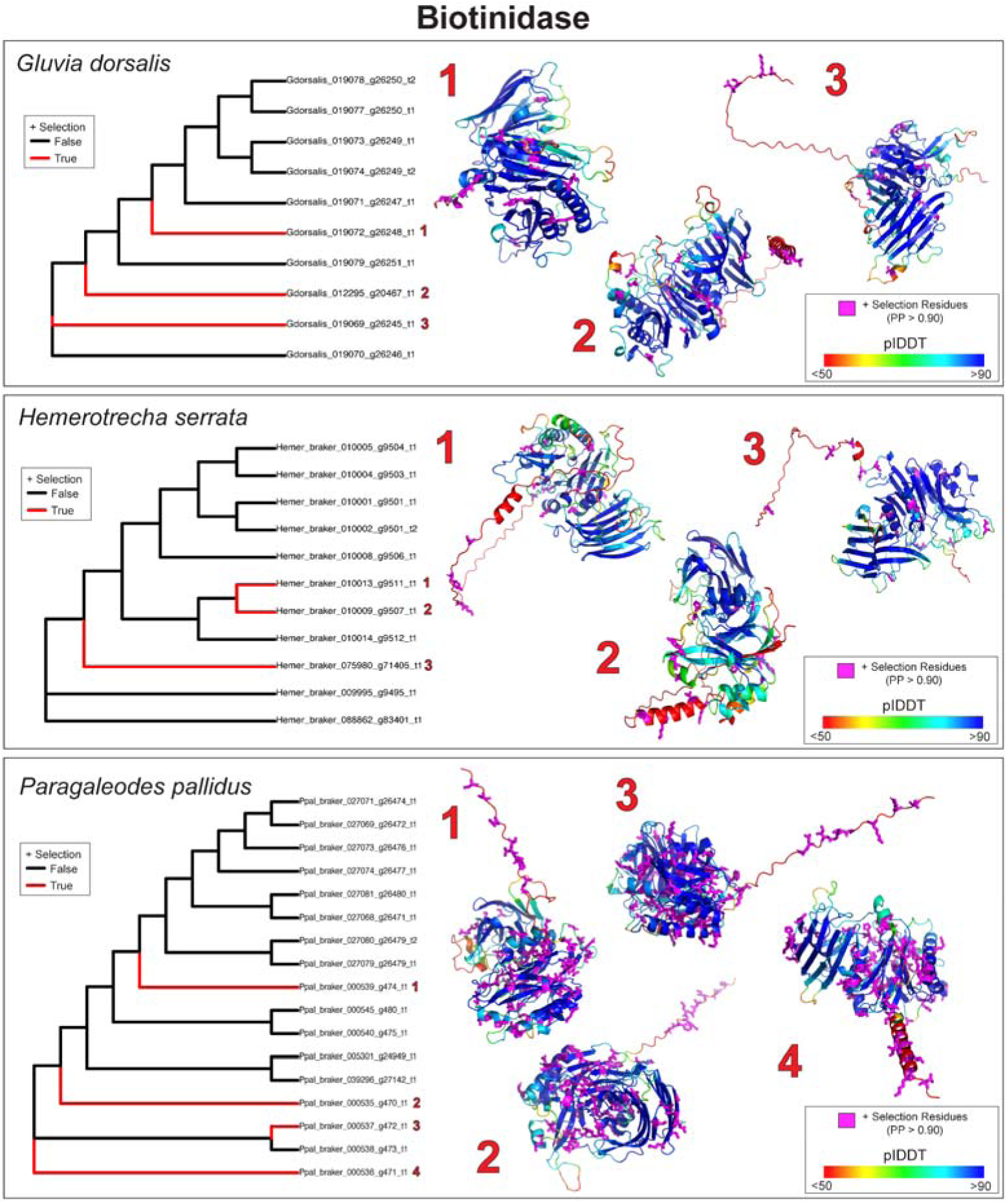
Inparalog positive selection results from aBSREL and corresponding 3D structural modeling of each respective positively selected protein for the functionally annotated Biotinidase-like protein (N0.HOG0000712) in solifuges. Maximum likelihood (ML) phylogenetic analysis and predicted 3D protein models for *Gluvia dorsalis* (top panel) and *Hemerotrecha serrata* (bottom panel). Left subplots display gene trees with all branches tested for episodic positive selection. Branches highlighted in red indicate inparalogs under significant positive selection (+ Selection (True) ; ω> 1, corrected p-value < 0.05), while black branches indicate no detected selection (False). Specific selected paralogs are numbered corresponding to their predicted structural models on the right. Structural models are colored by AlphaFold 3 per-residue confidence scores (pLDDT), ranging from very low confidence (red <50) to high confidence (blue >90). Pink coloration highlights the specific residues predicted to be under positive selection with a posterior probability (PP) > 0.90 recovered in FUBAR.

**Figure 5.**
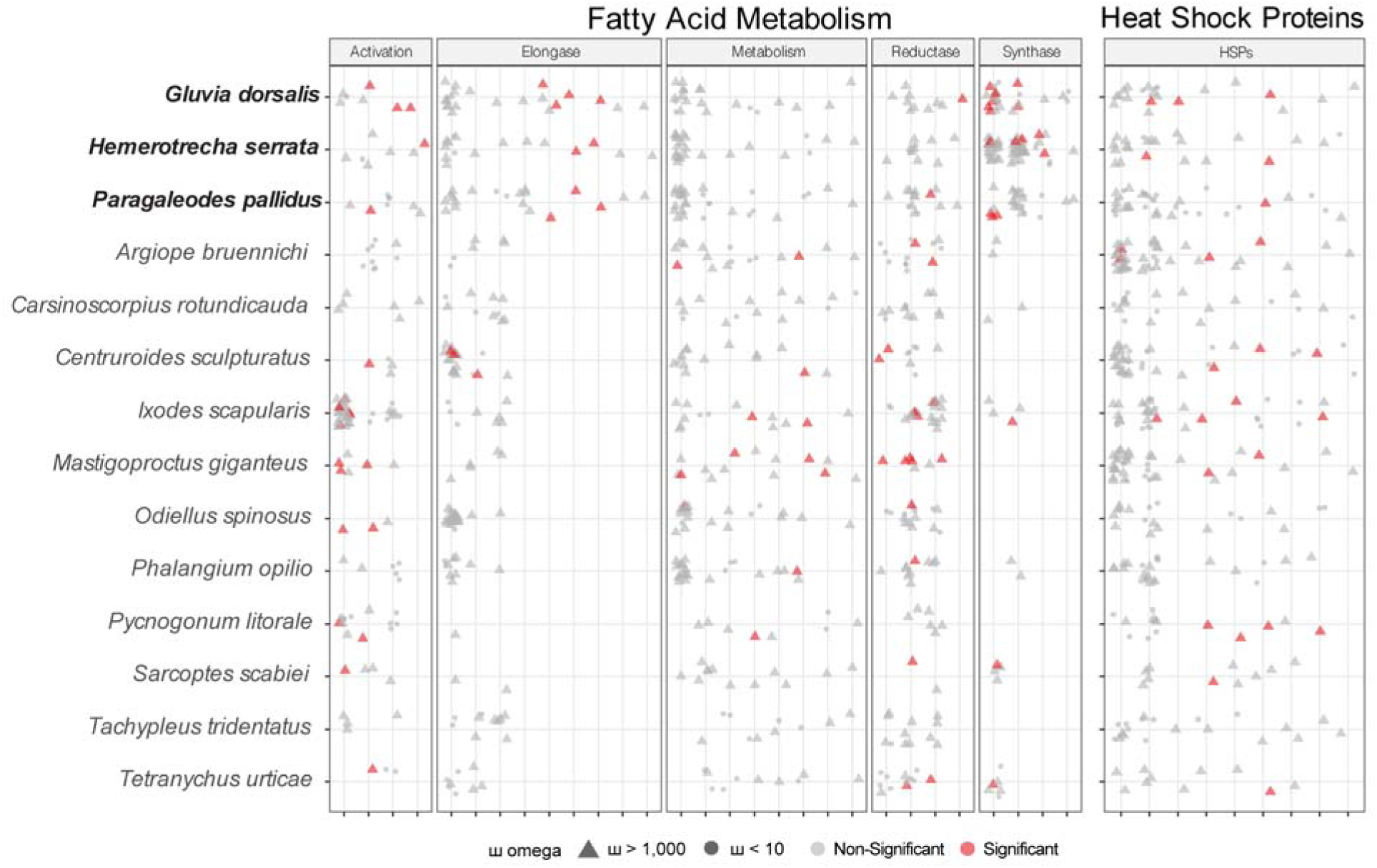
Comparative analysis of the evolutionary selection pressures on genes involved in Fatty Acid Metabolism and Heat Shock Proteins (HSPs) across various ingroup chelicerates. Solifugae species are species highlighted in bold. The columns categorize genes into functional pathways, split into two panels. The first panel denotes those Hierarchical Orthogroups (HOGs) involved in Fatty Acid Metabolism and is divided by Activation, Elongase, Metabolism, Reductase, and Synthase functional annotations. The second panel represents the HOGs associated with Heat Shock Proteins (HSPs). Individual data points represent one individual sequence assigned to a specific HOG depicted on the x-axis. Triangles represent sequences that recovered ω (omega) values greater than 1,000 in our aBRSEL analyses, while circles represent values < 10. Triangles in red denote positive selection (uncorrected p-value < 0.05), while triangles in grey represent those with uncorrected p-values > 0.05).

We delved into the 17 expanded HOGs demonstrating episodic advantageous selection to determine which site-specific residues test may have historically experienced constant selection. Proteins were functionally annotated as the cytochrome P450 family 2 (CYP2) featured the highest number of amino acid residues under positive selection (Supporting Information 3). This finding strongly underscores the CYP2 family as a key candidate gene family for future research that is unique to the solifuge lineage. Interestingly, three HOGs, despite presenting evidence for episodic selection, failed to recover amino acid residues under selection (Table 1). This result suggests that these specific in-paralogs perhaps gave these species a selective advantage for a short period of time. The functional annotation of these HOGs most closely resembled SideStep proteins (N0.HOG0000025), THAP domain-containing protein 2 (N0.HOG00001033), and were associated with serine hydrolase activity (N0.HOG0003779). The remaining paralogs examined all exhibited at least one amino acid residue that experienced constant positive selection.

We found that many of the site-specific residues in each of those sequences that underwent episodic selection are predominantly hydrophobic residues, which may suggest that they played pivotal roles in both providing structural stability to a protein by safeguarding the exposure of hydrophobic residues to the solvent (Xu et al., 1997; Okada et al., 2010; Gromiha et al., 2013) or to increase the catalytic capacity of an enzyme by helping to lower the activation energy required for a catalytic reaction (Bartlett et al., 2002). The bonds formed by hydrophobic residues has been indicated to strengthen with increasing temperatures, thus is one of the several contributing factors that strengthen the overall stability of a protein (Scheraga et al., 1962) Additionally, due to their hydrophobic nature, protein transmembrane domains are primarily hydrophobic, however some of the hydrophilic residues play crucial roles in stabilizing protein complexes (Dawson et al., 2003), while also facilitating effective receptor binding (Freeman-Cook et al., 2005). Although the difference in the number of hydrophobic and hydrophilic residues was minor, the selection pressure favoring hydrophobic residues suggests they may play a key role in enhancing protein stability, perhaps in warmer temperatures, or for boosting the catalytic activity of an enzyme.

Although functional protein research in arachnids remains heavily skewed toward mite, spider, and scorpion taxa, we herein summarize the broader functional implications of the advantageous paralogs identified within our solifuge dataset. While further empirical research is required to elucidate the precise role of each sequence, these findings establish a foundational baseline for future solifuge proteomics.

### Shared evidence of positive selection of inparalogs within expanded orthogroups among ingroup solifuges

#### Cytochrome P450 family 2

The cytochrome P450 superfamily are among the largest gene families responsible for encoding enzymes and are widely present in bacteria (Munro & Lindsay, 1996), plants (Nelson et al., 2008), and humans (Gonzalez, 1992). These enzymes are recognized for their diverse functional repertoire, which ranges from metabolizing a various endogenous and exogenous compounds (Chang & Kam, 1999), to synthesizing of ecdysone in insects, an important modulator in molting and metamorphosis (Rewitz et al., 2007). Additionally, they have been implicated in suppressing metabolic rates during periods of starvation in spiders (G. Zhang et al., 2025) and are sometimes expressed at high temperatures (Liu et al., 2017). In our positive selection analyses, all ingroup solifuges recovered at least one paralog within the cytochrome P450 family 2 classification (CYP2), and we recovered two separate expanded HOGs that closely matched CYP2. When focusing on arachnid CYP2 genes, a study on a predatory mite recovered one CYP2 ortholog related to ecdysteroid biosynthesis (Daimon et al., 2012; Wu & Hoy, 2016a). The remaining CYP2 genes identified lacked strong orthology to other arthropods and thus may be due to recent gene expansion events (Wu & Hoy, 2016), or may be unique to chelicerates. Because CYP2 genes have been implicated in metabolic-rate suppression, an especially relevant function for desert arachnids that may endure prolonged fasting, and in responses to elevated temperatures, their presence in our arid-adapted solifuge exemplars raises the possibility that this gene family contributes to desert adaptation in sarachnids.

The pattern of CYP major family expansion among arachnids appears to be consistent, at least among scorpions (Cao et al., 2013), spiders (G. Zhang et al., 2025), and mites (Grbić et al., 2011; Bajda et al., 2015; Shi et al., 2015; Wu & Hoy, 2016a). Considering the genomic evidence rendered from three solifuges, this appears to be true for the Solifugae. However, our analyses indicate that only a small subset of these genes are under selection (Table 1). All three solifuges possessed at least three positively selected paralogs assigned to the CYP2 gene family (Table 1; Figure 3), with an additional few more paralogs maintained in *Gluvia dorsalis* and *Hemerotrecha serrata* belonging to a completely separate HOG (Table 1, Supplemental Figure S2). Among those paralogs, in the HOG containing all Solifugae represented for in-paralog selection, many of these advantageous residues were concentrated at the core of the protein, with a few located near the N-terminus (Figure 3, Supplementary Information 2). The second CYP2 HOG supported more residues under positive selection, with an apparent mix of surface and interior residues (Supplemental Figure S2). Collectively, the two independent, CYP2-associated HOGs contain 20–40 gene copies for Solifugae–a range comparable to that observed in mites (Bajda et al., 2015; Wu & Hoy, 2016). While the drivers behind the higher number of positively selected transcripts in *Hemerotrecha serrata* (n=5) and *Gluvia dorsalis* (n=5) compared to *Paragaleodes pallidus* (n=3), remain unclear, the CYP2 clan offers a compelling target for further research into gene family expansion dynamics and functional roles within Solifugae.

### Biotinidase-related/Vanin family

The Vanin (VNN) gene family is a family of proteins that share considerable central domain similarities, which some researchers suggest that these proteins in the family, like Biotinidase, all bear enzyme activity (Granjeaud et al., 1999). Pantetheine hydrolases, or vanins, are conserved enzymes that hydrolyze pantetheine to yield pantothenic acid (vitamin B5), a critical precursor for the biosynthesis of coenzyme A (Naquet et al., 2014; Logan-Garbisch et al., 2015), which plays a critical role in energy production and fatty acid synthesis (Barritt et al., 2024). Extensive research regarding the functional role of these enzymes have primarily focused on humans and other mammals (Jansen et al., 2009; Kaskow et al., 2012; Bartucci et al., 2019), yet within these systems, evidence indicates that not only are they involved in coenzyme A synthesis, but also in inflammatory responses, oxidative stress, cell adhesion, and metabolism regulation (Bartucci et al., 2019). Moreover, mammalian pantetheine hydrolases show remarkable resistance to chemical and high heat exposure, comparable to thermophilic enzymes sourced from thermophilic organisms (Pitari et al., 1996). Among the few available terrestrial arthropod studies, it has been demonstrated that there is an upregulation of genes encoding for pantetheine hydrolases after induced oxidative stress in *Drosophila* larvae (Logan-Garbisch et al., 2015), are downregulated in low temperatures in the lepidopteran fall armyworm (Vatanparast & Park, 2022), and play a role in viral transmission in ticks and mosquitoes (Grabowski et al., 2017, 2019; Rezende et al., 2024). Despite the limited research on VNN family proteins in arthropods, the evolutionary implications are profound given their role in metabolic flexibility (e.g. fatty acid synthesis, energy resources) and general immunity (e.g. antioxidant activity).

Within our solifuge ingroup, all three species exhibited three or more positively selected amino acid sequences containing Biotinidase/VNN family signatures (Table 1; Figure 4). Our results recovered comparatively fewer residues under positive selection for *Gluvia dorsalis* and *Hemerotrecha serrata*, whereas amino acid positively selected residues for *Paragaleodes pallidus* were prolific throughout the tertiary protein structure (Figure 4; Supporting Information 3). In all three cases, several advantageous residues were localized near the N-terminus, which may suggest that this protein evolved effective cell signaling (Varland et al., 2015). For *P. pallidus*, on the other hand, this enzyme may have evolved additional advantages, such as for promoting structural integrity or for engaging efficient catalytic activity necessary for success in new environments.

While the precise functional role of these proteins remains poorly characterized in arthropods, and arachnids specifically, their conservation and positive selection result suggests a significant adaptive advantage in solifuges among the expanded orthogroup compared to the other chelicerates examined. These proteins may enhance solifuge fitness by modulating metabolic pathways, potentially supporting fatty acid synthesis, meeting high locomotor energy demands, or enhancing cellular resilience to oxidative stress. By optimizing these physiological mechanisms, such adaptations would possibly enable arid-adapted solifuges to withstand prolonged periods of starvation by efficiently utilizing lipid stores and oxidative stress caused by extreme thermal stress typical of desert environments.

### Lineage-specific and parallel in-paralog positive selection within expanded solifuge orthogroups

### Side-step proteins

One of the few characterized orthogroups shared between all ingroup solifuges were amino acid sequences with an affinity for sidestep (Side) proteins, which are muscle-derived transmembrane proteins that are essential for motor axons to enter muscle targets, and essentially increase muscle attractiveness for axon growth (Sink et al., 2001; Kinold et al., 2021; Heymann et al., 2022). In conjunction with the Beaten Path (Beat), proteins expressed on motor axons, the interaction of the two immunoglobulin proteins are necessary to begin the formation of synapses (Siebert et al., 2009). Studies demonstrate that mutations in the Side protein cause dramatic locomotion defects and irreparable misallocation of nerves in *Drosophila* larvae (Sink et al., 2001; Kinold et al., 2021). No orthologs of Beat and Side have been identified in vertebrates (Cortés et al., 2023), yet eight or more paralogous copies of Beat and Side, respectively, have been revealed in *Drosophila* (Cortés et al., 2023; Osaka et al., 2024). Of the limited studies on Beat and Side, they have solely focused on elucidating their exact function in the *Drosophila* model system (Sink et al., 2001; Siebert et al., 2009; Kinold et al., 2021; Heymann et al., 2022; Osaka et al., 2024). Few of those copies are associated with locomotion and leg innervation, yet the functions of other paralogs in the central nervous system have yet to be elucidated.

Given that solifuges have amino acid sequences that most closely match Side proteins, this suggests that Side proteins may be largely conserved across distant invertebrates, but have only been thoroughly studied in *Drosophila.* Curiously, only *H. serrata* and *P. pallidus* were recovered to have two amino acid sequences under strong positive selection with affiliation to Side proteins (Table 1); both are members of the suborder Boreosolifugae and inhabit hotter and more xeric desert climates than G. dorsalis, which occupies more Mediterranean dry-summer, semiarid systems.

As Solifugae are capable of remarkable running speeds, the inference of positive selection for proteins related to locomotion is consistent with their physiology. The implications of Side proteins under positive selection suggests that this lineage of solifuges may have undergone selective pressure for producing efficient motor control to accommodate for specialized, fast movements and/or to increase process signaling of motor neural networks. Curiously, however, *P. pallidus* did not recover any residues supporting positive selection signal and our analyses only recovered four residues with signal (Supplemental Figure S4; Supporting Information 3). This result suggests that these proteins may no longer be experiencing persistent selection, but these proteins facilitated their success in their present environment. Nevertheless, beyond existing studies on the mechanics of speed of some of the fastest arthropods (Bartholomew et al., 1985; Pfeffer et al., 2019), many of which are adapted to desert and semi-desert environments, exploring how Side influences arthropod velocity may highlight additional functional importance of these proteins in arthropods.

### Alpha-Tocopherol transfer protein/phosphatidylinositol bisphosphate (PIP2) binding

Among the many physiological responses of heat-induced stress, oxidative stress leads to an increase in reactive oxygen species (ROS), which in turn, causes significant lipid damage (peroxidation), protein degradation, and DNA degradation (Schieber & Chandel, 2014). While low concentrations of ROS may play important roles in several biological processes, such as cell signaling (Suzuki et al., 1997; Gabbita et al., 2000) or transcription regulation (Siauciunaite et al., 2019; Hong et al., 2024), biological mechanisms are necessary to restore homeostasis during elevated ROS levels to ensure an organism’s survival. An intrinsic defense mechanism that many organisms possess is the presence of antioxidative enzymes and molecular antioxidants (Felton & Summers, 1995; Zhao et al., 2020a; Nie et al., 2023) which are a line of protection against highly reactive ROS that jeopardize the configuration of biomolecules and threaten their normal functionality. Such antioxidant responses, in addition to the production of heat shock proteins, have been widely considered to be important factors for heat tolerance among insects (Jena et al., 2013; Ju et al., 2014; Bianchini et al., 2016; Z. Q. Miao et al., 2020; González-Tokman et al., 2025), and with fewer studies recognizing its importance in arachnids (Zhao et al., 2020b; Fu et al., 2022; Nie et al., 2023). The consistent prevalence of these complex defense mechanisms across distant organisms suggests that these defense strategies are ancient and were perhaps crucial physiological adaptations to mitigate the effects of temperature stress in terrestrial ecosystems.

The results of our paralog selection analysis showed evidence that *P. pallidus* was the only solifuge species to have two amino acid sequences with affinities to alpha-tocopherol transfer proteins (α-TTPs) and phosphatidylinositol biphosphate (PIP_2_) binding that were under positive selection (Table 1; Supplemental Figure S4). The number of advantageous residues were notable (40), which their distribution on the protein primarily near alpha-helices, with some near the N-terminus (Supplemental Figure S4). First, α-TTP are cystosolic, lipid-binding proteins that facilitate the secretion of α-tocopherol (i.e. vitamin E), an antioxidant, across a lipid membrane (Catignani, 1975; Panagabko et al., 2003). At least in mammals, it has been demonstrated that α-TTP exhibits a high binding affinity to α-tocopherol compared to other lipophilic molecules, and can selectively uptake α-tocopherol compared to other tocopherol isoforms (Panagabko et al., 2003). Moreover, molecular simulations of α-TTP binding with different tocopherol isoforms reinforces a high ligand specificity to α-tocopherol as the binding to other isoforms may cause instability in the post-bounded molecular complex (Helbling et al., 2014). In addition to α-tocopherol binding, it has been demonstrated that α-TTP can directly bind to phosphatidylinositol phosphates (PIPs), which not only promotes the release of α-tocopherol (Lamprakis et al., 2015), but their position within the membrane are key landmarks for the anchoring of α-TTP (van den Bogaart et al., 2011; Lamprakis et al., 2015). With regard to *P. pallidus,* the specimen was collected from Egypt, whose geographical position is recognized as part of the Saharan desert (Walter & Breckle, 1986). This area of vast and extreme desert habitat holds one of the hottest annual maximum land surface temperatures in the world (Mildrexler et al., 2006; Hu et al., 2023), thus carrying implications regarding the positive selection signal recovered by our aBRSEL analyses for *P. pallidus.* Few arachnid studies have observed an increase in gene expression levels of antioxidant enzyme-coding genes in response to oxidative stress, as these are common biomarkers to measure stress (Chainy et al., 2016; Yousef et al., 2017; Aziz et al., 2020).

However, despite changes in antioxidant enzyme levels in heat (Zhao et al., 2020b), or toxic metal environments (Aziz et al., 2020), the role of antioxidant transfer proteins and their relationship to heat stress is unknown in arachnids. The selective advantage of α-tocopherol transfer-related proteins in *P. pallidus* suggests that the optimal binding of specific ligands, such as antioxidants or membrane lipids, may be a key adaptation for hot desert survival. Future studies employing comparative gene knockdowns of antioxidant transfer proteins and enzymes would illuminate the mechanisms used to withstand oxidative stress induced by extreme heat.

### Gene expression modifiers: Transposons, Transposases, Helix domains, and Transcription factors

Transposable elements (TEs), or transposons, often encode enzymes (e.g. transposases) that facilitate their integration into genomes (Kojima, 2019), and in some cases, these elements are fully autonomous, harboring the specialized machinery for their own transposition (Cohen & Shapiro, 1980).

In eukaryotes, they occur in considerable proportions of the genome (Kidwell, 2002; Petersen et al., 2019) and in chelicerate genomes in particular, most are DNA transposons with high average TE superfamily diversity (Petersen et al., 2019). Moreover, TEs are significant sources of rapid genomic innovation and TE activity has been associated with the rise of novel traits promoting adaptive radiations (Feiner, 2016; Schrader & Schmitz, 2019; Carleton et al., 2020). In recent years, research has illuminated the diverse suite of TE functionalities, highlighting their complex contributions to an organism’s genomic architecture, such as their roles in genome reorganization (Kidwell, 2002; Alzohairy et al., 2013; Kojima, 2019), gene regulation (Bourque, 2009), and how some are activated by environmental stress stimuli (Casacuberta & González, 2013). Arthropod studies show that environmental stress can alter the activity of specific transposable element families, including under heat, oxidative, and starvation stress in *Drosophila* (Bodelón et al., 2023; Milyaeva et al., 2023; Merenciano et al., 2025). In arid habitats, where organisms like camel spiders repeatedly face thermal stress, oxidative stress, water limitation, and prolonged fasting, stress-responsive TEs may therefore represent an additional source of regulatory and genomic variation available to selection. Signatures of positive selection of TEs or transposases would suggest that they may have been co-opted for functional roles in an organism’s genome. Conversely, signatures of negative or purifying selection suggest that genomic sequences are under evolutionary constraint to preserve their original function.

Our aBSREL analyses revealed signatures of positive selection in transposon/transposase-annotated sequences in *H. serrata* and *G. dorsalis*. Notably, *H. serrata* yielded four sequences from four distinct expanded orthogroups with evidence of episodic selection, whereas only a single sequence was recovered for *G. dorsalis* (Table 1; Supplemental Figure S5A&D). This sequence belonged to a shared HOG with *H. serrata* (N0.HOG0000947), however for *H. serrata* our aBSREL results supported ∼60% of sites in *H. serrata* evolved under positive episodic selection, with the remaining sites exhibited purifying selection (ω = 0; Supporting Information 2). A similar pattern was detected in a second, independently annotated transposon in *H. serrata* (N0.HOG0000311), with approximately 40% of sites showing signatures of episodic selection, and the remaining 60% subject to purifying selection. Our FUBAR analyses, on the other hand, detected fewer residues under selection for N0.HOG0000947 for both species (Supplemental Figure S5D; Supporting Information 3) and for *H. serrata* (N0.HOG0000823; Supplemental Figure S5C; Table 2). However, no evidence supported any residues under positive selection for *H. serrata* N0.HOG0001033 (Supplemental Figure S5A). This result suggests these particular genes may have offered a fitness advantage shortly after duplication (Ohta, 1994; Kondrashov et al., 2002), for a brief period, and that only a few amino acids, if any, continued to provide a selective advantage. While the sites under purifying selection support the inference that certain regions remain essential for function, the significant proportion of sites under positive selection highlights a strong potential for the emergence of novel functionalities. These include evolving preferences for integration in different target areas of the genome (Cosby et al., 2019), increasing enzyme flexibility associated with DNA binding (Majumdar & Rio, 2015; Ghanim et al., 2019), or evading host silencing by producing mutations that leave no identifiable sequence for silencing (Cosby et al., 2019; Lawlor & Ellison, 2023).

Although the exact role of these transposons/transposases are unknown in solifuges (as in other chelicerate groups), in other arthropod systems, TEs can be inserted into a gene transcript to provide a selective advantage by influencing melanization in moths (Hof et al., 2016), promoting insecticidal resistance (Salces-Ortiz et al., 2020; Zidi et al., 2022), mitigating oxidative stress (Merenciano et al., 2016), or leveraging adaptive success in new environments (González et al., 2009, 2010). Many TEs, however, are also subject to high mutation rates (Ho et al., 2021) or may accumulate mutations upon being silenced, thereby subjecting them to the loss of the ability to transpose (Betancourt et al., 2024). In the curious case of *H. serrata,* four independent sequences were found to be under selection, suggesting that TEs play an important role in their biology, in contrast to *G. dorsalis* and *P. pallidus*. Future studies regarding the timing of TE emergence and location could unlock essential implications about adaptive evolution in solifuges.

Although TEs are a prominent source for modulating gene expression in eukaryotes, partially serving as transcription factor binding sites (Trizzino et al., 2017; Panigrahi & O’Malley, 2021; Gebrie, 2023), many zinc finger proteins also function as transcription factors that recognize specific DNA sequences, yet their overall functional capacity is diverse (Laity et al., 2001). Similarly, helix domains are common DNA-binding motifs that are key protein domains for transcription regulation, yet in eukaryotes they also play a role in protein-protein interactions or protein signaling (Aravind et al., 2005). We recovered sequences for *Pa. pallidus* and *G. dorsalis* with signatures of positive selection for a functionally annotated Helix-turn-Helix (HTH) domain and a zinc finger protein (ZFP), respectively (Table 1; Supplemental Figure S5B&F). Both HTH domains and ZFPs are small structural motifs involved in gene expression (Pellegrini-Calace & Thornton, 2005; Vilas et al., 2018). However, ZFPs are critical stress-responsive proteins that prevent protein denaturation and degradation, thereby play a critical role in driving high temperature tolerance (Miao et al., 2025). Conversely, HTHs have been implicated in mitigating abiotic stress, specifically in drought, in predominantly plant systems (X. Li et al., 2006; Govind et al., 2009; Castilhos et al., 2014; Zuo et al., 2023). While, to our knowledge, HTHs have yet been empirically linked to stress tolerance in arthropods, evidence suggests that transcription factors play a role in detoxification during heavy metal exposure in spiders (J. Wang et al., 2021). Aligned with these findings, our positive selection results for *Pa. pallidus* and *G.dorsalis* suggests that perhaps transcription factors play an essential role in mediating adaptive responses to arid and semi-arid environments.

### Immunoglobulin-like domains and metabolism

Immune defense in vertebrates is characterized as an adaptive immune system involving a large repertoire of antigen-specific receptors achieved through somatic gene rearrangement and clonal selection at the genomic level (Papavasiliou et al., 1997; D. Jung et al., 2006). To date, no antibodies have been detected in invertebrates, although immunoglobulin-like domains appear to be a common and conserved feature (Lanz Mendoza & Faye, 1999; Mandrioli et al. 2015), many of which play an important role in immune response (Jung et al., 2019), or in the nervous system (Yoshihara et al., 1991; Bothwell, 2006; H. Li et al., 2025). However, members of the immunoglobulin superfamily (IgSF), have been linked to other roles such as in cell-to-cell adhesion, recognition, and signal transduction (H. Li et al., 2025). A recent study examined the diversification of the IgSF Down syndrome cell adhesion molecule (Dscam) and revealed that Chelicerata evolved a conserved, yet lineage-specific repertoire of nonclassical Dscams (H. Li et al., 2025). Moreover, many of these Dscams are structurally and mechanistically distinct from insects (Yue et al., 2016), thus presenting an intriguing focus for future study.

The results of our analyses indicate that *H. serrata* and *P. pallidus* both possess at least one immunoglobulin-like domain under positive episodic selection (Table 1; Supplemental Figure S6A &D), with fewer than 10 residues demonstrating evidence for positive selection (Supplementary Information 2. Since immunoglobulin-like domains can be found in diverse proteins, and through gene duplication and later divergence in chelicerates, it is difficult to link these domains to a specific functional role at this time. However, due to their general conservation and their role within other arthropod groups, these domains are likely to be associated with a neuronal or immune function (Yue et al., 2016; H. Li et al., 2025). As part of the IgSF superfamily, Sidestep proteins were also curiously limited to *H. serrata and P. pallidus,* and we therefore speculate that these specific immunoglobulin-domains could be playing a neuronal-related fitness advantage possibly associated with their mobility.

Long-chain fatty acid-CoA ligases (ACSLs) play crucial intermediary roles in fatty acid metabolism by activating free long-chain fatty acids to form acyl-CoA in the fatty acid biosynthesis pathway for storage, lipid membrane biosynthesis, or for energy production (Steinberg et al., 2000; Digel et al., 2009). In arthropods, ACSL has been documented to play a key role in lipid accumulation, however their isoforms can be expressed in different tissues, and may be highly expressed during pre- or early diapause (C. Zhang et al., 2019; Xiang et al., 2021), or during prolonged starvation (Alves-Bezerra et al., 2016). However, the main metabolic fate of a long-chain fatty acid is through oxidation to render ATP (Digel et al., 2009), in which an ACSL must activate the fatty acid prior to it being transported across an intracellular membrane for further energetic processing (Watkins, 1997). ACSL enzyme isoforms are known to maintain different substrate specificities that preferentially activate specific fatty acids (Fujino et al., 1996; Kang et al., 1997; Jepson et al., 2014), and the overexpression of ACSL isoforms results in enhanced fatty acid uptake (Digel et al., 2009). Some evidence suggests that a subcellular spatial organization of ACSL activity in different membranes (e.g. endoplasmic reticulum, mitochondrial, plasma) is present for different ASCL enzymes, all of which may be designated for a specific metabolic pathway (Lewin et al., 2001; Digel et al., 2009; O’Sullivan et al., 2012).

*H. serrata* was the only species to recover an ACSL paralog under episodic selection (Table 1; Supplemental Figure S6B), and only one positively selected residue was recovered from our FUBAR analysis. This latter result indicates that this sequence was historically important for adaptive success, yet subsequently underwent fixation due to the lack of amino acid residues under positive selection (Kosiol et al., 2008). Since the primary outcome of a long-chain fatty acid is for energy production, we hypothesize that this specific paralog may have been retained to provide this species with an energy-related fitness advantage, possibly related to enhanced muscle function since this species also recovered other muscle-related functional paralogs under selection (Table 1). The fitness advantage of this ACSL may have evolved an alternative capacity to activate a diverse array of fatty acids for use in energetically demanding processes, allowing for faster and more efficient loading in the binding site for activation. Alternatively, this paralog may be associated with motor axon bundles as documented in *Drosophila* (O’Sullivan et al., 2012), which may also be related to muscle function. A promising avenue for future investigation would be to narrow down which tissues express this paralog to elucidate its functional significance.

Lastly, we recovered one paralog for *P. pallidus* that matched neuroligin and bile salt-activated ligase (Table 1; Supplemental Figure S6C), both members belonging to the α//3-hydrolase fold superfamily, under strong positive episodic selection (Chatonnet et al., 2017a) with four sites under directional selection (Supporting Information 3). However, it is worth noting that invertebrates lack conventional bile salts (Reschly et al., 2008), therefore a bile salt associated function can be eliminated as a possible advantageous role in solifuges. The α//3-hydrolase superfamily, along with other well-known protein superfamilies (e.g. cytochrome P450 (CYP) superfamily), has been documented to also be functionally diverse ranging from lipases, acetylcholine esterases, proteases, reductases, and more (Ollis et al., 1992). Despite their widely recognized role in catalytic activity, some proteins belonging to this superfamily, like neuroligin, lack enzymatic activity altogether (Krejci et al., 1991; Lenfant et al., 2014). Neuroligin homologues have been detected in several invertebrate species and have been implicated in sensory or synaptic modulation (Biswas et al., 2008; Wu & Hoy, 2016b; Corthals et al., 2017; Durand et al., 2021), as they are primarily expressed in pre- and post- synaptic membranes (Knight et al., 2011).

Nevertheless, members within this superfamily have undergone significant gene expansions in animals, and some proteins have acquired novel functions unrelated to their associated role in processes like neurotransmission (Chatonnet et al., 2017b). Since studies on arachnid proteins pertaining to this specific family are rare, and lipases have been historically annotated according to mammalian homologues (Chatonnet et al., 2017b), it is difficult to formulate hypotheses regarding the functional significance of this protein family. However, due to their generally conserved structure, it is likely that this protein may be a lipase specific to solifuges since many proteins in the genome are associated with lipid production (see below) or may be involved in neural activity.

### Serine hydrolase activity

In addition to the VNN gene family, serine hydrolases are a large family of functional enzyme classes that are present in all forms of life with a conserved catalytic mechanism (Kumar et al., 2021), thus reinforcing the importance of this gene family in critical fundamental biological processes. In characterizing serine hydrolases, it appears that the number of overall nonredundant active serine hydrolases are the highest present in the adult stage in *Drosophila* (Kumar et al., 2021). One of the key characteristic roles that these enzymes play is their role in lipid metabolism, however expression profiling studies of this superfamily suggest that these genes are expressed in specific tissues (Kumar et al., 2021; Z. Miao et al., 2023). However, in the tobacco hornworm, these enzymes have other functional roles in indigestion, detoxification, neurotransmission, among other biological functions (Miao et al., 2023).

While functional roles of serine hydrolase enzymes are versatile and broad, the general categories of functionality can be deduced into few enzyme categories that may be of importance to the two solifuge species from the Old World, *G. dorsalis* and *P. pallidus*. Curiously, no selection signal was detected for *H. serrata* (Table 1) and no advantageous residues were identified for *G.dorsalis* (Supplemental Figure S7). The precise drivers associated with possible implications for a positive selection signal in those two species are unclear, however, we suspect that the multiple copies in *G. dorsalis* may possess functional differences that have promoted their success.

### Fatty acid synthesis and metabolism

Solifuges have undoubtedly undergone significant evolutionary changes that have drastically changed their lipid profile compared to other chelicerate groups (Figure 5), specifically among their fatty acid elongases (ELOs) and fatty acid synthases (FASs). While both enzymes are involved in one of the many assembly lines of broad fatty acid metabolism, ELOs are crucial in regulating the length of fatty acid chains (D.-T. Li et al., 2019), and FASs are involved in the biosynthesis and storage of lipids (Maier et al., 2010). In arthropods, ELOs have been implicated largely in their role in the production of cuticular hydrocarbons (CHCs; Chung et al., 2014; Falcón et al., 2014; Finck et al., 2016; D.-T. Li et al., 2019; Pei et al., 2021), which play a vital role in desiccation resistance. Beyond their role in cuticular waterproofing, ELOs also involved in pheromone biosynthesis (Jurenka, 2004; Wicker-Thomas et al., 2015; Luo et al., 2025) and these two pathways require a very long-chain fatty acids as a precursor (Stanely-Samuelson et al. 1988). In generating a very long-chain fatty acid, an FAS is a key enzyme that continues a multi-step cycle for building a long fatty acid chain. Among some of the documented terrestrial arthropod examples such as in dipterans (Stanely-Samuelson et al. 1988), lepidopterans (Lambremont, 1971), and hypothesized to be the case in at least one mite (Brückner & Heethoff, 2020), evidence suggests that the principal length of fatty acids that are synthesized by arthropods FASs are approximately 16 to 18 carbon atom fatty acid chains. Fatty acids of this length are classified as long-chain fatty acids, of which are primarily destined for metabolic energy (Digel et al., 2009). The production of CHCs, wax esters, or pheromones requires a very-long fatty acid chain as a precursor (Stanley-Samuelson et al., 1988). Very-long fatty acid chains are generally classified as fatty acid chains with > 22 carbon atoms and to achieve this length, an ELO is required to elongate a long-chain fatty acid (Kaczmarek & Boguś, 2021). Moreover, elongases have been implicated in enhancing desiccation resistance in Sonoran desert *Drosophila* in gene knockout experiments (Z. Wang et al., 2023), therefore we suspect that elongases may be playing a similar crucial role in solifuges desert habitats where they have evolved a novel repertoire of elongases primarily to alter the cuticular membrane composition for desiccation resistance.

One possibility is that synthases in solifuges may have evolved novel solutions to minimize water loss, as well as meet the high-intensity metabolic demands during their periods of activity and for reproduction. First, there is a general consensus that long-distance migratory insects use lipids as the dominant energy source for sustained flight (van der Horst et al., 1993; Haunerland, 1997; X. Li et al., 2023). To support the remarkable endurance and speed documented in solifuges (Muma, 1966b; Wharton, 1986; Punzo, 1998b), a taxon documented to sustain this behavior over several hours, an internal mechanism must exist to rapidly mobilize energy. We suspect that lipids are the underlying fuel for this sustained activity, typical of such highly mobile arachnids. To meet the high-intensity metabolic demands during any endurance exercise, taxa that engage in this type of behavior have not only remodeled their fatty acid (e.g. lipid) capacity, but this is often coupled with an elevated antioxidant capacity to mitigate oxidative damage (McWilliams & Muller, 2026), which is also evidenced by this study in all three species (e.g. Biotinidase).

Although all three solifuges appear to uniquely possess multiple synthases under positive selection compared to our other examined chelicerate groups, only one HOG was shared among all three taxa (Figure 5). Curiously, *H. serrata* and *G. dorsalis* share an additional HOG with affiliation to FASs, and *H. serrata* exclusively possesses a third synthase-related HOG (Figure 5). Because FASs catalyze the *de novo* synthesis of long-chain fatty acids, the positive selection detected in the two additional HOGs for our female solifuge species may indicate a selective advantage to directly benefit oocyte maturation or the high energy demands required to prepare for and sustain embryonic development, ultimately increasing fecundity in harsh environments. In other arthropod groups, while FAS is expressed in both males and females, FAS expression is highest in females (J.-Y. Cheng et al., 2023; Xin et al., 2025) and in ovarian tissues (J.-Y. Cheng et al., 2023). Moreover, several FAS gene silencing experiments across several arthropod groups have been shown to negatively impact fecundity (Alabaster et al., 2011; L. Li et al., 2016; J.-Y. Cheng et al., 2023; Xin et al., 2025),with some evidence directly linking an impairment of both oocyte development and ovary maturation (Hu et al., 2026). We hypothesize that these noted differences may be fatty acid composition differences between male and female solifuges. Since the physiological demands for reproduction are energetically costly during vitellogenesis and oocyte growth, a significant amount of lipid storage is crucial to ensure reproductive success (Lease & Wolf, 2011; Fruttero et al., 2017; Romero et al., 2018; Garcia et al., 2025). In several studies on reproductive behavior in solifuges, many female solifuge species are known to retreat into burrows for egg deposition (Punzo 1998b; Muma, 1966c; Wharton, 1986; Hrušková-Martišová et al., 2007, 2010b; Erdek & Kayhan, 2016), and in at least one species, refraining from feeding a few days prior to oviposition (Erdek & Kayhan, 2016). Burrowing behavior for reproduction is hypothesized to be family-specific (Erdek & Kayhan, 2016); some behavioral data suggest that it may endure between half to one hour (Muma, 1966c; Wharton, 1986), suggesting that this behavior requires high metabolic endurance to achieve desired burrow depths. Moreover, females have been observed to deposit at least two non-consecutive egg clutches, with oviposition lasting approximately 60-90 minutes, and females subsequently refrain from feeding for a significant period post-oviposition (Muma, 1966c; Erdek & Kayhan, 2016). Considering these stages from pre- to post-oviposition and observed maternal care of oocytes, reproduction is a highly demanding process that undoubtedly requires a significant amount of energy. We suspect that the role of FAS in solifuges not only independently evolved to change the molecular architecture of the solifuge cuticle and for meeting intense exercise demands, but also to give solifuges a selective advantage for successful reproduction in xeric environments.

### Heat shock proteins

Heat shock proteins (hsps) belong to a broader family of molecular chaperones, proteins that perform essential cellular maintenance functions, including preventing aberrant protein aggregation and facilitating the refolding of denatured proteins to their native conformations (reviewed in Sørensen et al., 2003; Evgen’ev et al., 2015; Chen et al., 2018). As a result, hsps are central mediators of the cellular stress response, and their expression is strongly induced by a range of environmental stressors, most notably heat stress (King & MacRae, 2015). Notably, hsp genes, in particular those from the hsp70 family, are among the most evolutionary conserved genes known (Villar, 2000). For example, humans and *Drosophila* hsp70 share 73% amino acid sequence identity (Hunt & Morimoto, 1985). This high degree of conservation reflects their role across the tree of life, including in organisms exposed to extreme thermal environments. For example, desert ants (*Cataglyphis bombycina*) experience body temperatures exceeding 50°C, with their hsp70 levels elevated even prior to heat exposure, suggesting a preemptive thermoprotective mechanism (Wehner et al., 1992; Gehring & Wehner, 1995). Further, variation in hsps gene copy number has been documented across a wide range of taxa and is frequently interpreted as an adaptive response to thermal stress (e.g., Kim et al., 2022; Gordillo-Perez et al., 2026). In this study, we found no evidence of hsp gene family expansion in any of the three solifuge species, suggesting that adaptation to desert conditions in these arachnids has not involved changes in hsp gene repertoire. Nevertheless, molecular evolution analyses revealed signatures of positive selection in one hsp HOG across all three species, with two additional HOGs under positive selection in *G. dorsalis* and one further HOG in *H. serrata* (Figure 5). Together, these findings suggest that while hsp gene number remained stable in Solifugae, sequence-level adaptive evolution has occurred in a subset of these genes. This may point to a potential role for expression level of hsps as adaptations to desert thermal regimes. Pairing these new genome assemblies with RNAseq data and physiological experiments may therefore be a highly valuable future avenue of investigation in solifuge biology.

## Conclusion

Despite the renowned physiological capabilities of Solifugae and their tendency among arachnids to occupy arid and semiarid habitats, the molecular underpinnings of their adaptations to these environments are virtually unknown. Through an exploratory comparative genomic analysis, we identified expanded orthogroups specific to the order Solifugae and evaluated selective pressures acting upon paralogous sequences of expanded gene families. Our analysis revealed significant signatures of positive selection within several key gene families across the order, many of which have plausible links to survival in desert and semidesert environments. Notably, paralogous sequences associated with the cytochrome P450 and biotinidase gene families exhibited consistent evidence of selection among all three taxa, suggesting that metabolic regulation, detoxification, oxidative stress mitigation, and nutrient processing may represent important components of solifuge adaptation to resource limited and thermally challenging habitats. Other shared candidate loci under selection were implicated in axonal guidance and lipid metabolism, potentially linking genomic evolution to the remarkable locomotive capacity and energetic demands of these animals. Together, these results suggest that solifuge genomes contain a diverse molecular toolkit for coping with the physiological challenges of arid environments. The identification of these candidate loci provides a preliminary genetic foundation for understanding the selective advantages that have shaped solifuge evolution and spotlights priorities for future investigations of functional genomics in this unusual arachnid order.

## Acknowledgements

We are grateful to the University of Wisconsin-Madison for providing Center for High Throughput Computing (CHTC) resources. This work was supported by the National Science Foundation (NSF) through the NSF PRFB Award Number 2305969 to E.L.G, NSF grant no. IOS-2016141 to P.P.S, and NSF grant DEB-1754030 to M.R.G. Additional funding was provided by the CSU AAUP grant X31083 to C.E.S.L. We thank Gustavo Hormiga for providing access to SK to the Pegasus High Performance Computing cluster at George Washington University for assembly and annotation purposes. SK was supported by the ANRF Ramanujan Fellowship (RJF/2023/000045).

## Notes

### Competing Interest Statement

The authors have declared no competing interest.

