## Supplemental Material for "Comparative genomics of the unusual arachnid order Solifugae spotlight the molecular and genetic basis for adaptations to arid habitats"

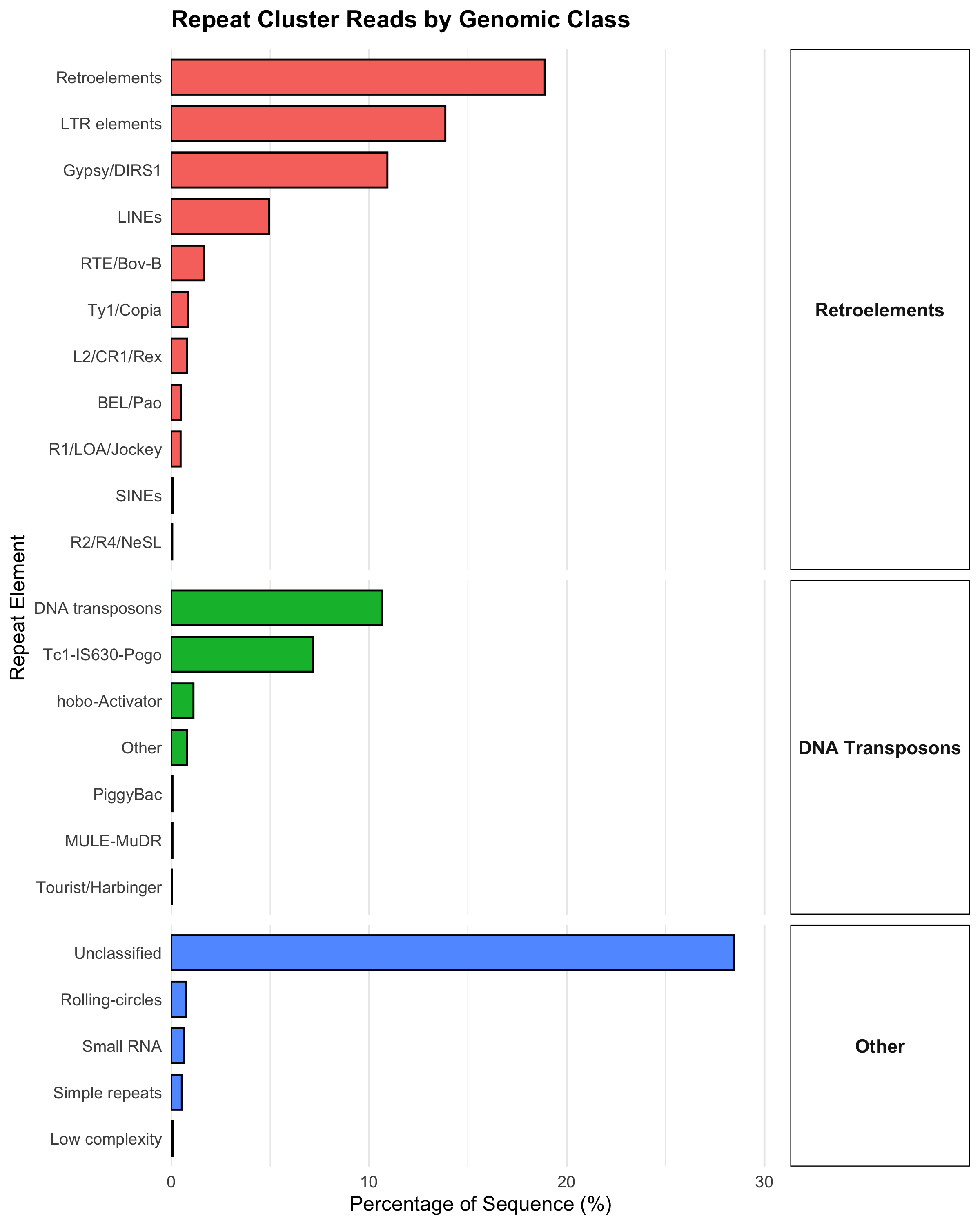

**Figure 1. Genomic landscape classifications of repeat elements in *Hemerotrecha serrata* assembly from RepeatMasker analysis.** Bar chart summarizes the percentage of the total sequences by identified repeat categories and is partitioned hierarchical classification into three distinct genomic categories: Retroelements (top panel), DNA Transposons (middle panel), and Other genomic feature types (bottom panel).

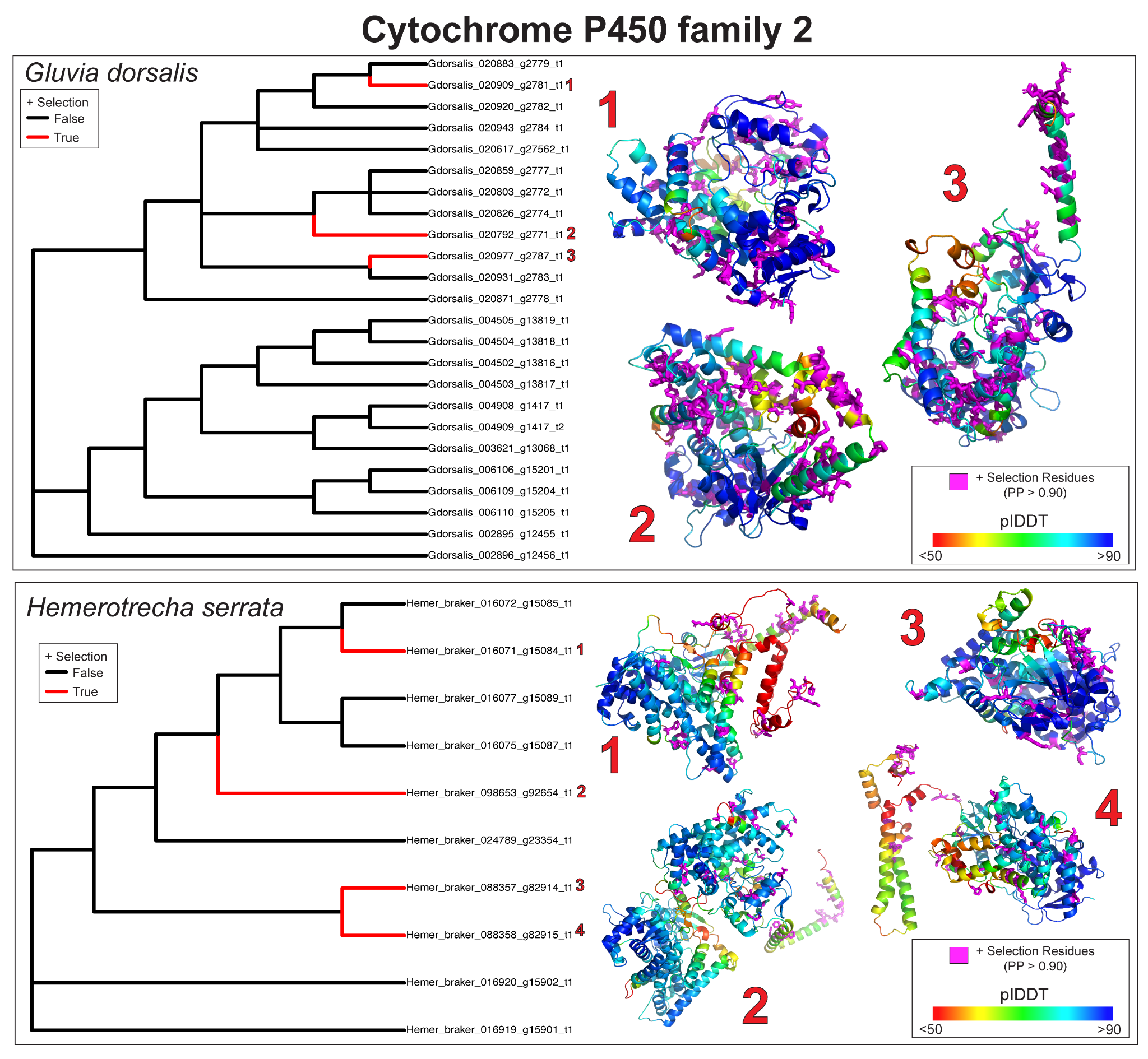

**Figure 2. Inparalog positive selection results from aBSREL and corresponding 3D structural modeling of each respective positively selected protein for the functionally annotated Cytochrome P450 family 2 protein (N0.HOG0000708) in solifuges.** Maximum likelihood (ML) phylogenetic analysis and predicted 3D protein models for *Gluvia dorsalis* (top panel) and *Hemerotrecha serrata* (bottom panel). Left subplots display gene trees with all branches tested for episodic positive selection. Branches highlighted in red indicate inparalogs under significant positive selection (+ Selection (True) ; ω> 1, corrected p-value < 0.05), while black branches indicate no detected selection (False). Specific selected paralogs are numbered corresponding to their predicted structural models on the right. Structural models are colored by AlphaFold 3 per-residue confidence scores (pLDDT), ranging from very low confidence (red <50) to high confidence (blue >90). Pink coloration highlights the specific residues predicted to be under positive selection with a posterior probability (PP) > 0.90 recovered in FUBAR.

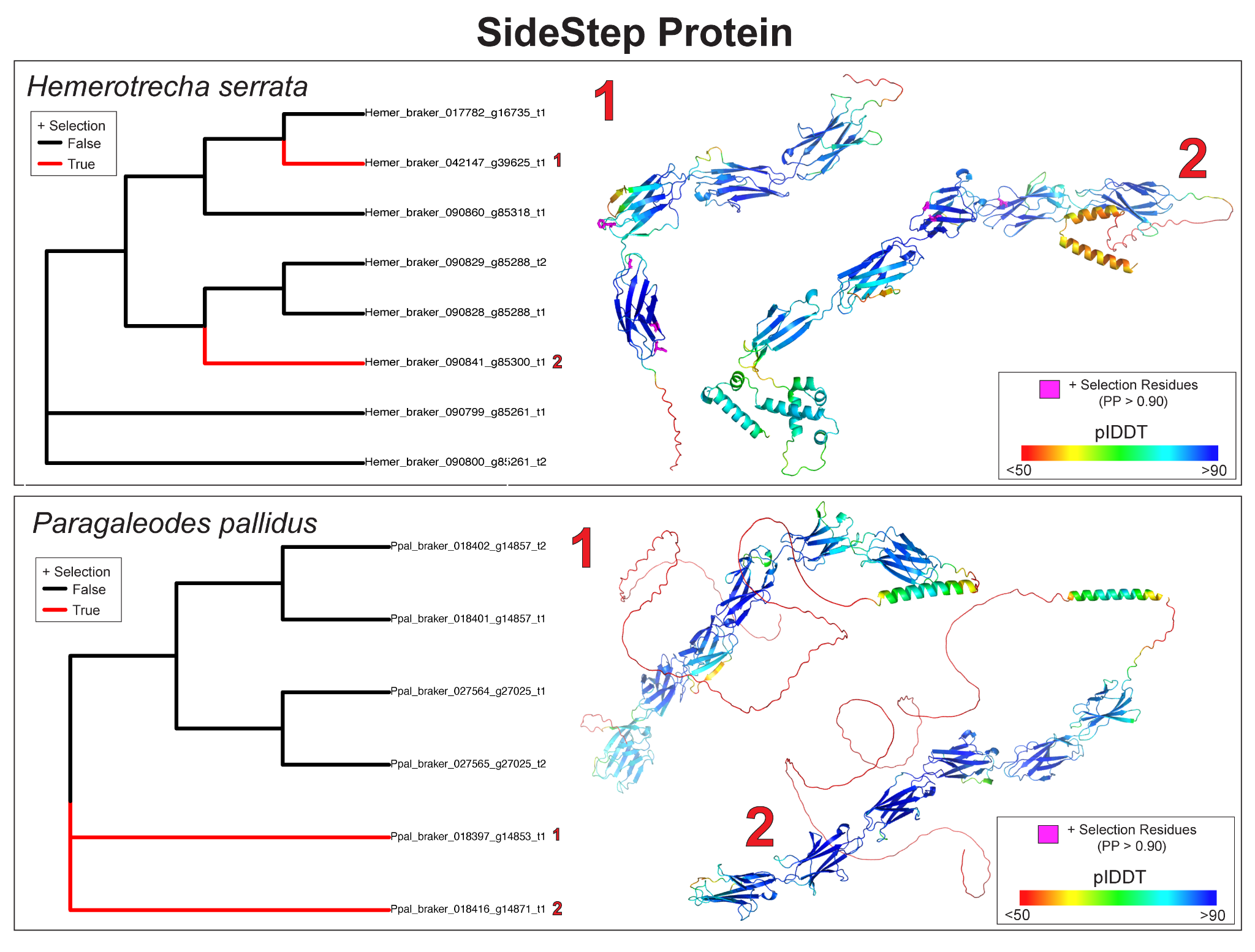

**Figure 3. Inparalog positive selection results from aBSREL and corresponding 3D structural modeling of each respective positively selected protein for the functionally annotated SideStep protein (N0.HOG 0000025) in solifuges.** Maximum likelihood (ML) phylogenetic analysis and predicted 3D protein models for *Hemerotrecha serrata* (top panel) and *Paragaleodes pallidus* (bottom panel). Left subplots display gene trees with all branches tested for episodic positive selection. Branches highlighted in red indicate inparalogs under significant positive selection (+ Selection (True) ; ω> 1, corrected p-value < 0.05), while black branches indicate no detected selection (False). Specific selected paralogs are numbered corresponding to their predicted structural models on the right. Structural models are colored by AlphaFold 3 per-residue confidence scores (pLDDT), ranging from very low confidence (red <50) to high confidence (blue >90). Pink coloration highlights the specific residues predicted to be under positive selection with a posterior probability (PP) > 0.90 recovered in FUBAR.

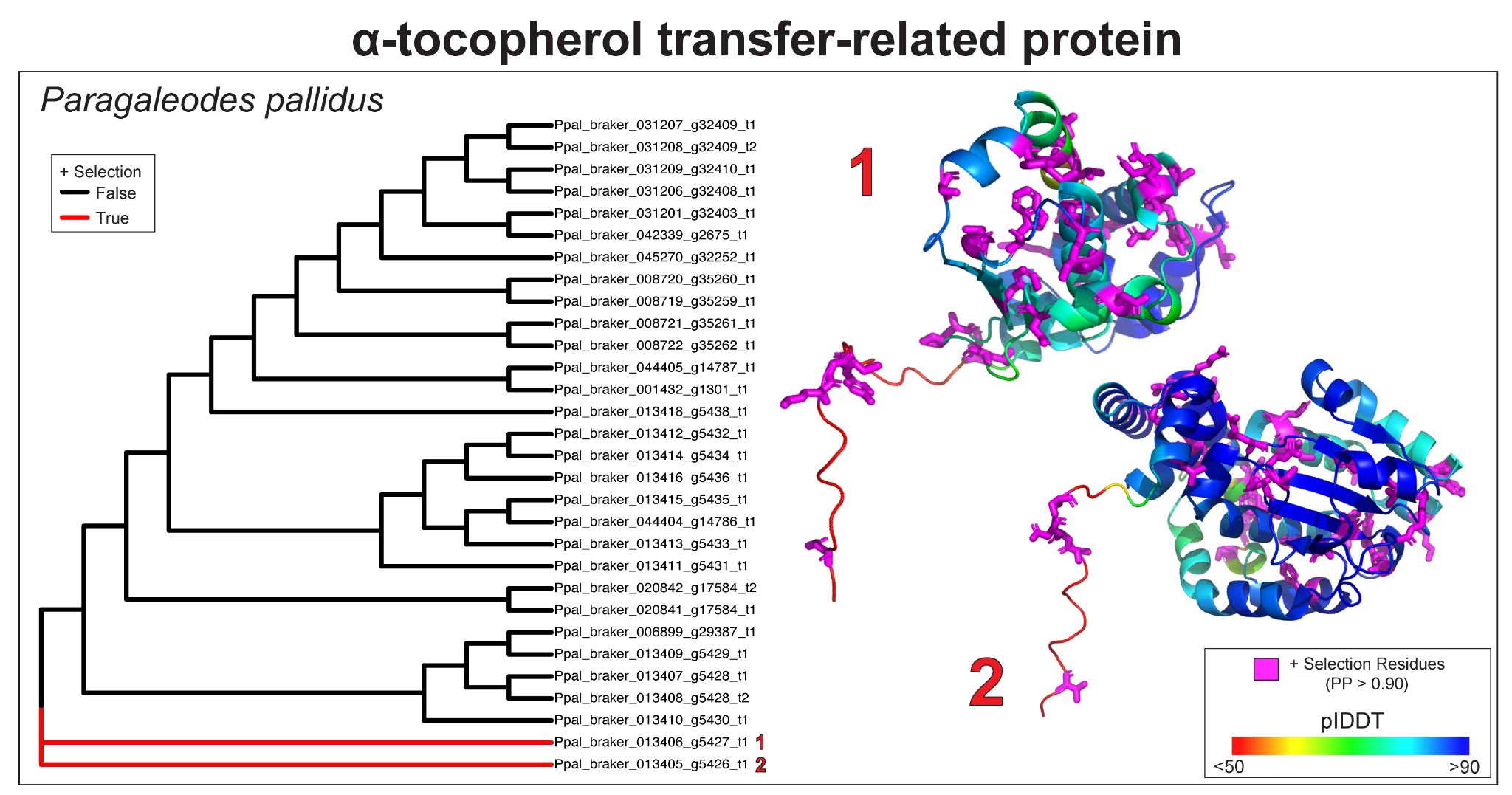

**Figure 4. Inparalog positive selection results from aBSREL and corresponding 3D structural modeling of each respective positively selected protein for the functionally annotated SideStep protein (N0.HOG0000154) in solifuges.** Maximum likelihood (ML) phylogenetic analysis and predicted 3D protein models for *Paragaleodes pallidus*. Left subplots display gene trees with all branches tested for episodic positive selection. Branches highlighted in red indicate inparalogs under significant positive selection (+ Selection (True) ; ω> 1, corrected p-value < 0.05), while black branches indicate no detected selection (False). Specific selected paralogs are numbered corresponding to their predicted structural models on the right. Structural models are colored by AlphaFold 3 per-residue confidence scores (pLDDT), ranging from very low confidence (red <50) to high confidence (blue >90). Pink coloration highlights the specific residues predicted to be under positive selection with a posterior probability (PP) > 0.90 recovered in FUBAR.

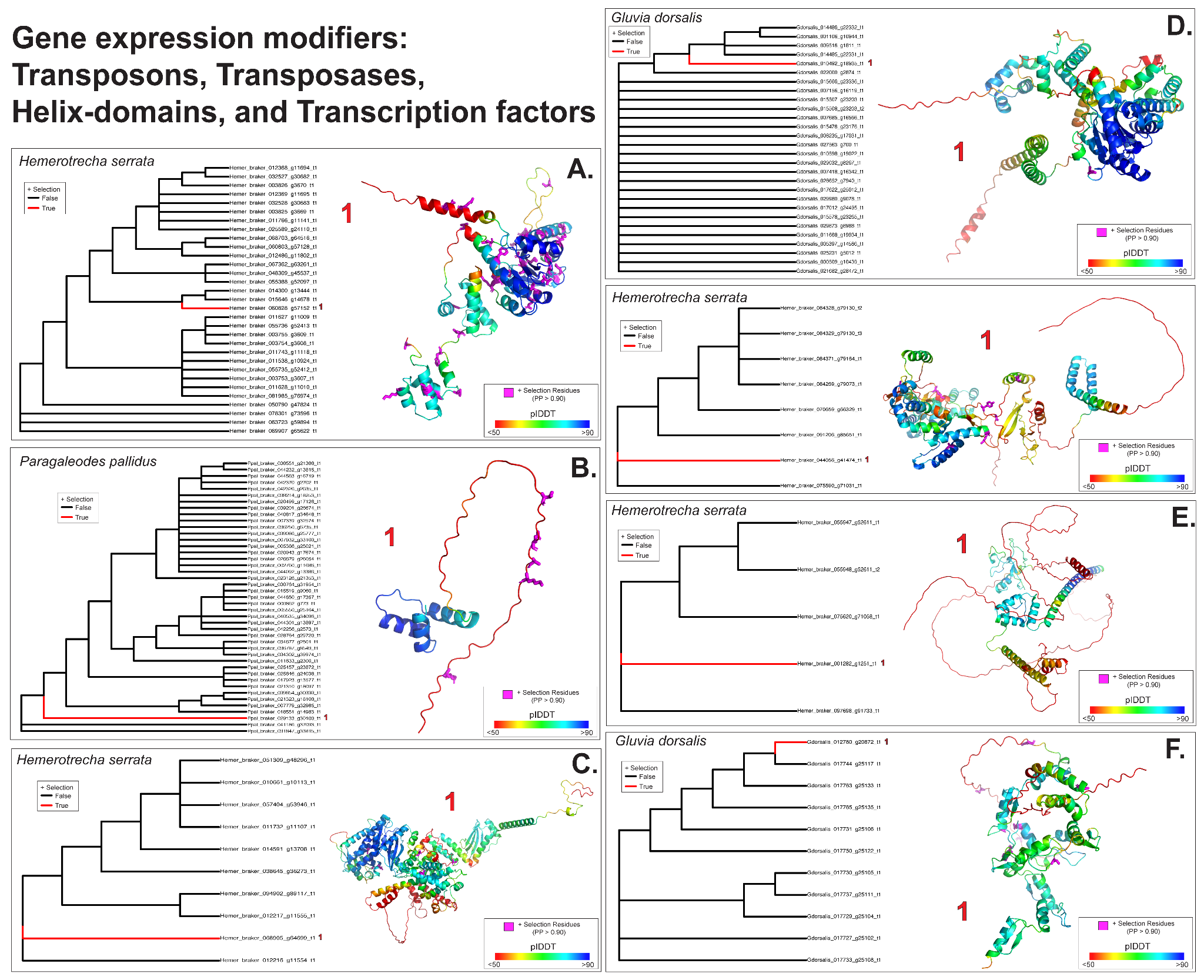

**Figure 5. Inparalog positive selection results from aBSREL and corresponding 3D structural modeling of each respective positively selected protein for the functionally annotated proteins broadly assigned to gene expression modifiers in solifuges.** All panels (A-F) contain maximum likelihood (ML) phylogenetic analysis and predicted 3D protein models. Left subplots display gene trees with all branches tested for episodic positive selection. Branches highlighted in red indicate inparalogs under significant positive selection (+ Selection (True) ; ω> 1, corrected p-value < 0.05), while black branches indicate no detected selection (False). Specific selected paralogs are numbered corresponding to their predicted structural models on the right. Structural models are colored by AlphaFold 3 per-residue confidence scores (pLDDT), ranging from very low confidence (red <50) to high confidence (blue >90). Pink coloration highlights the specific residues predicted to be under positive selection with a posterior probability (PP) > 0.90 recovered in FUBAR. A.) Selection results of N0.HOG0000311 for *Hemerotrecha serrata* transposon. B.) Results of N0.HOG0000470 for *Paragaleodes pallidus* helix-turn helix domain C.) Results of N0.HOG0000823 for *Hemerotrecha serrata* transposase (TNP-like RNase H N-terminal domain). D.) Results of N0.HOG0000947 for *Gluvia dorsalis* and *Hemerotrecha serrata* transposon. E.) Results of N0.HOG0001033 *Hemerotrecha serrata* THAP domain-containing protein. F.) Results of N0.HOG0003779 for *Gluvia dorsalis* zinc finger protein.

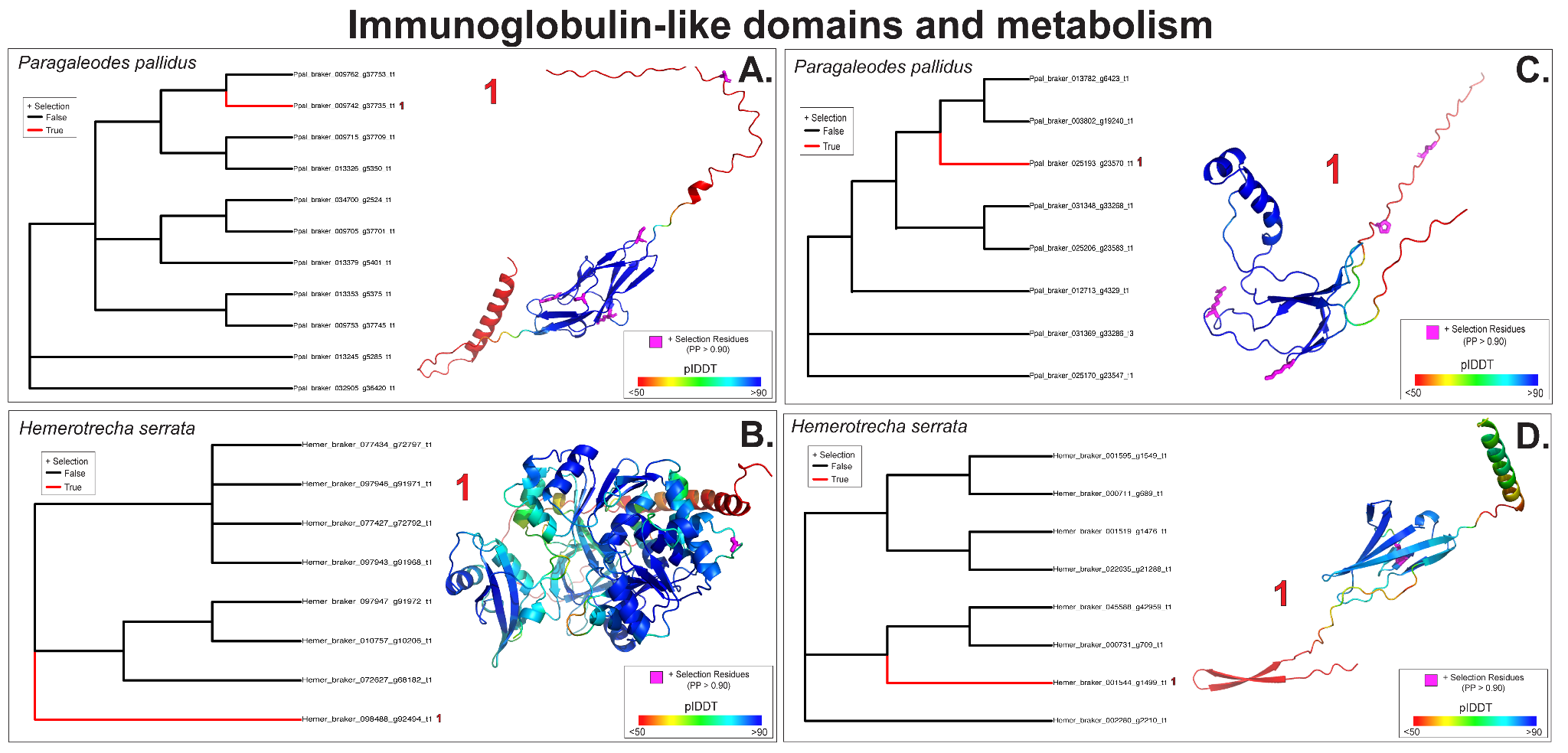

**Figure 6. Lineage-specific inparalog positive selection results from aBSREL and corresponding 3D structural modeling of each respective positively selected protein for the functionally annotated proteins broadly assigned to immunoglobulins-like proteins or metabolism in solifuges.** All panels (A-F) contain maximum likelihood (ML) phylogenetic analysis and predicted 3D protein models. Left subplots display gene trees with all branches tested for episodic positive selection. Branches highlighted in red indicate inparalogs under significant positive selection (+ Selection (True) ; ω> 1, corrected p-value < 0.05), while black branches indicate no detected selection (False). Specific selected paralogs are numbered corresponding to their predicted structural models on the right. Structural models are colored by AlphaFold 3 per-residue confidence scores (pLDDT), ranging from very low confidence (red <50) to high confidence (blue >90). Pink coloration highlights the specific residues predicted to be under positive selection with a posterior probability (PP) > 0.90 recovered in FUBAR. A.) Selection results of N0.HOG0000448 for *Paragaleodes pallidus* immunoglobulin-like protein. B.) Results of N0.HOG0000636 for *Hemerotrecha serrata* for long-chain fatty acid-CoA ligase C.) Results of N0.HOG0000914 for *Paragaleodes pallidus* neuroligin and bile salt activated lipase. D.) Results of N0.HOG0001409 for *Hemerotrecha serrata* immunoglobulin.

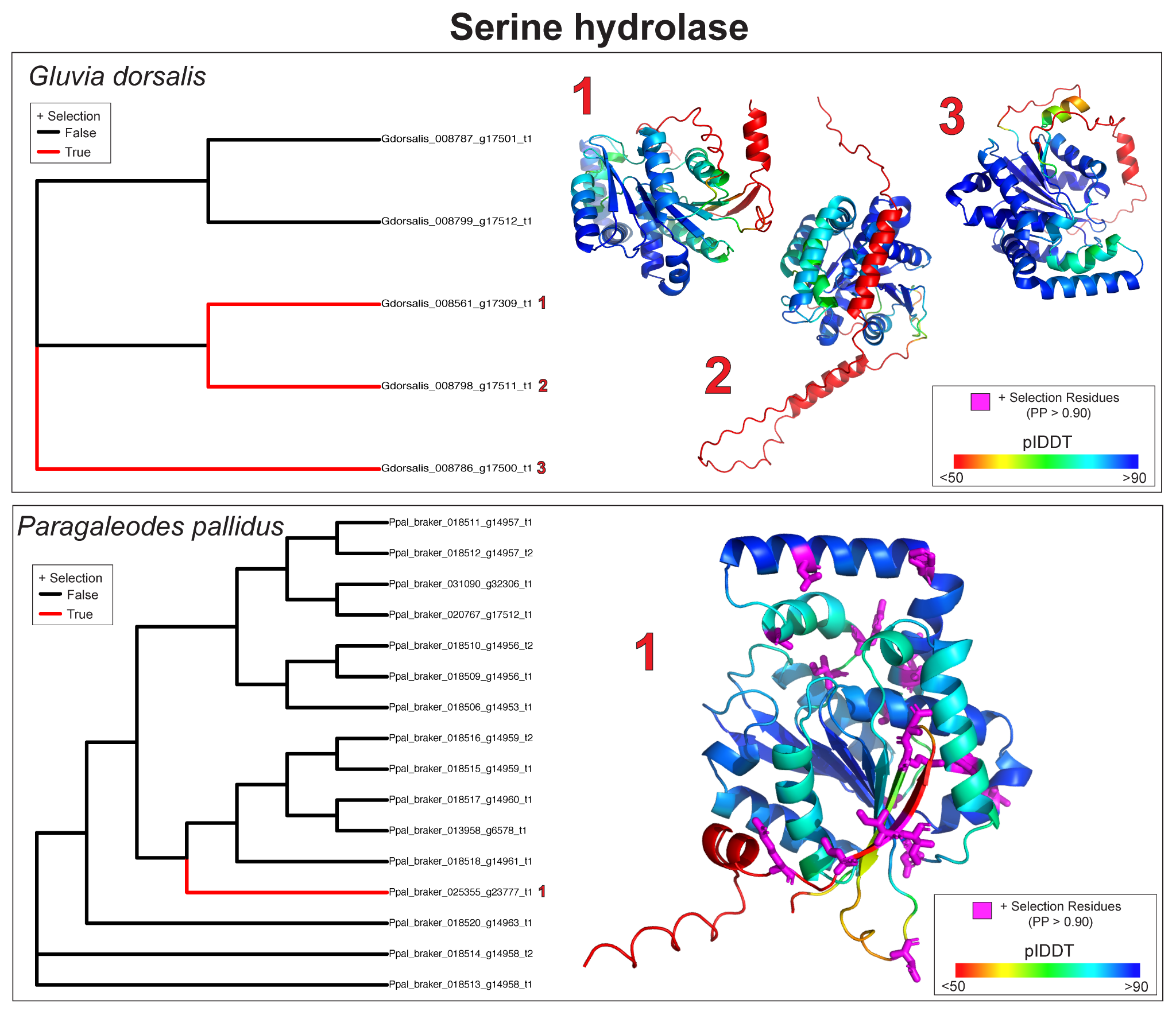

**Figure 7. Inparalog positive selection results from aBSREL and corresponding 3D structural modeling of each respective positively selected protein for the functionally annotated serine hydrolase protein (N0.HOG0001600) in *Gluvia dorsalis* and *Paragaleodes pallidus*.** Maximum likelihood (ML) phylogenetic analysis and predicted 3D protein models for *Gluvia dorsalis* (top panel) and *Paragaleodes pallidus* (bottom panel).Left subplots display gene trees with all branches tested for episodic positive selection. Branches highlighted in red indicate inparalogs under significant positive selection (+ Selection (True) ; ω> 1, corrected p-value < 0.05), while black branches indicate no detected selection (False). Specific selected paralogs are numbered corresponding to their predicted structural models on the right. Structural models are colored by AlphaFold 3 per-residue confidence scores (pLDDT), ranging from very low confidence (red <50) to high confidence (blue >90). Pink coloration highlights the specific residues predicted to be under positive selection with a posterior probability (PP) > 0.90 recovered in FUBAR.
